# Loss of centromere function drives karyotype evolution in closely related *Malassezia* species

**DOI:** 10.1101/533794

**Authors:** Sundar Ram Sankaranarayanan, Giuseppe Ianiri, Marco A. Coelho, Md. Hashim Reza, Bhagya C. Thimmappa, Promit Ganguly, Rakesh Netha Vadnala, Sheng Sun, Rahul Siddharthan, Christian Tellgren-Roth, Thomas L Dawson, Joseph Heitman, Kaustuv Sanyal

## Abstract

Genomic rearrangements associated with speciation often result in chromosome number variation among closely related species. *Malassezia* species show variable karyotypes ranging between 6 and 9 chromosomes. Here, we experimentally identified all 8 centromeres in *M. sympodialis* as 3 to 5 kb long kinetochore-bound regions spanning an AT-rich core and depleted of the canonical histone H3. Centromeres of similar sequence features were identified as CENP-A-rich regions in *Malassezia furfur* with 7 chromosomes, and histone H3 depleted regions in *Malassezia slooffiae* and *Malassezia globosa* with 9 chromosomes each. Analysis of synteny conservation across centromeres with newly generated chromosome-level genome assemblies suggests two distinct mechanisms of chromosome number reduction from an inferred 9-chromosome ancestral state: (a) chromosome breakage followed by loss of centromere DNA and (b) centromere inactivation accompanied by changes in DNA sequence following chromosome-chromosome fusion. We propose AT-rich centromeres drive karyotype diversity in the *Malassezia* species complex through breakage and inactivation.

## Introduction

Centromeres are the genomic loci on which the kinetochore, a multi-subunit complex, assembles to facilitate high fidelity chromosome segregation. The centromere-specific histone H3 variant CENP-A is the epigenetic hallmark of centromeres, as it replaces canonical histone H3 in the nucleosomes to make specialized centromeric chromatin that acts as the foundation to recruit other kinetochore proteins. A remarkable diversity in the organization of centromere DNA sequences has been observed to accomplish this conserved role (Roy and Sanyal, 2011, Yadav et al., 2018b).

The smallest known centromeres are the point centromeres present in budding yeasts of the family *Saccharomycetaceae* that span <200 bp in length (Clarke and Carbon, 1980, Gordon et al., 2011, Kobayashi et al., 2015). These centromeres are organized into conserved DNA elements I, II, and III that are recognized by a cognate kinetochore protein complex called the CBF3 complex, making them genetically defined centromeres. Small regional centromeres, identified in several *Candida* species, form the second category (Sanyal et al., 2004, Padmanabhan et al., 2008, Kapoor et al., 2015, Chatterjee et al., 2016) and have a 2 to 5 kb region enriched by kinetochore proteins. These centromeres can either have unique DNA sequences or a homogenized core that is flanked by inverted repeats. The third type of centromere structure is the large regional centromere, often repetitive in sequence and spanning over 15 kb. Large regional centromeres can be transposon enriched as in *Cryptococcus* species or organized into repeat structures around a central core as in *Schizosaccharomyces pombe* (Chikashige et al., 1989, Clarke and Baum, 1990, Sun et al., 2017, Yadav et al., 2018b).

While the organization of DNA elements is variable, a majority of known centromeres share AT-richness as a common feature. Examples include the CDEII of point centromeres, central core sequences in *S. pombe*, and centromeres of *Neurospora crassa*, *Magnaporthe oryzae*, *Plasmodium falciparum*, and diatoms (Fitzgerald-Hayes et al., 1982, Iwanaga et al., 2010, Rhind et al., 2011, Kapoor et al., 2015, Diner et al., 2017, Yadav et al., 2019). Even the recently described mosaic centromere structure observed in *Mucor circinelloides* that lost CENP-A comprises an AT-rich kinetochore-bound core region (Navarro-Mendoza et al., 2019). While suppression of recombination around centromeres has been correlated with reduced GC content (Lynch et al., 2010), the genetic underpinning of how an AT-rich DNA region favors kinetochore assembly remains unclear. Ironically, AT-rich sequences have been shown to be fragile sites of a chromosome (Zhang and Freudenreich, 2007).

Several lines of evidence suggest that centromeres are species-specific and are among the most rapidly evolving genomic regions, even between closely related species (Bensasson et al., 2008, Padmanabhan et al., 2008, Rhind et al., 2011, Roy and Sanyal, 2011). This is accompanied by the concomitant evolution of CENP-A and the associated kinetochore proteins (Talbert et al., 2004). Functional incompatibilities between centromeres result in uniparental genome elimination in interspecies hybrids (Ravi and Chan, 2010, Sanei et al., 2011). The divergent nature of centromeres is proposed to be a driving force for speciation (Henikoff et al., 2001, Malik and Henikoff, 2009).

Asexual organisms, by virtue of inter- and intra-chromosomal rearrangements, diversify into species clusters that are distinct in genotype and morphology (Barraclough et al., 2003). These genotypic differences include changes in both chromosomal organization and number. Centromere function is directly related to karyotype stabilization following a change in chromosome number. Rearrangements in the form of telomere-telomere fusions and nested chromosome insertions (NCIs), wherein an entire donor chromosome is ‘inserted’ into or near the centromere of a non-homologous recipient chromosome, are among the major sources of chromosome number reduction (Lysak, 2014). Such events often result in the formation of dicentric chromosomes that are subsequently stabilized by breakage-fusion-bridge cycles (McClintock, 1941) or via inactivation of one centromere through different mechanisms (Han et al., 2009, Sato et al., 2012). Well known examples of telomere-telomere fusions include the formation of extant human chromosome 2 by fusion of two ancestral chromosomes (Yunis and Prakash, 1982, Jdo et al., 1991), the reduction in karyotype observed within the members of the Saccharomycotina such as *Candida glabrata*, *Vanderwaltozyma polyspora*, *Kluyveromyces lactis,* and *Zygosaccharomyces rouxii* (Gordon et al., 2011), and the exchange of chromosomal arms seen in plants and fungi (Schubert and Lysak, 2011, Sun et al., 2017). NCIs have predominantly shaped karyotype evolution in grasses (Murat et al., 2010). Chromosome number reduction by centromere loss has also been reported (Gordon et al., 2011).

To investigate if centromere breakage can be a natural source of karyotype diversity in closely related species, we sought to identify centromeres in a group of *Malassezia* yeast species that exhibit chromosome number variation. *Malassezia* species are lipid-dependent basidiomycetous fungi that are naturally found as part of the animal skin microbiome (Theelen et al., 2018). The *Malassezia* genus presently includes 18 species divided into three clades – A, B, and C. These species also have unusually compact genomes of less than 9 Mb, organized into 6 to 9 chromosomes as revealed by electrophoretic karyotyping of some of these species (Boekhout and Bosboom, 1994, Boekhout et al., 1998, Wu et al., 2015). Fungemia-associated species like *Malassezia furfur* belong to Clade A. Clade B includes common inhabitants of human skin that are phylogenetically clustered into two subgroups namely Clade B1 that contains *Malassezia globosa* and *Malassezia restricta* and Clade B2 that contains *Malassezia sympodialis* and related species. Clade C includes *Malassezia slooffiae* and *Malassezia cuniculi* which diverged earlier from a *Malassezia* common ancestor (Wu et al., 2015, Lorch et al., 2018).

Besides humans, *Malassezia* species have been detected on the skin of animals. For example, *M. slooffiae* was isolated from cows and goats, *M. equina* from horses, *M. brasiliensis* and *M. psittaci* from parrots, and a cold-tolerant species *M. vesperilionis* isolated from bats (Lorch et al., 2018, Theelen et al., 2018). Additionally, culture-independent studies of fungi from environmental samples showed that the *Malassezia* species closely realted to those found on human skin were also detected in diverse niches such as deep-sea vents, soil invertebrates, hydrothermal vents, corals, and Antarctic soils (Amend, 2014). More than ten *Malassezia* species have been detected as a part of the human skin microbiome (Findley and Grice, 2014). The human skin commensals such as *M. globosa, M. restricta,* and *M. sympodialis* have been associated with dermatological conditions such as dandruff/seborrheic dermatitis, atopic dermatitis, and folliculitis (Theelen et al., 2018). Recent reports implicate *M. restricta* in conditions such as Crohn’s disease and *M. globosa* in the progression of pancreatic cancer (pancreatic ductal adenocarcinoma) (Aykut et al., 2019, Limon et al., 2019). Elevated levels of *Malassezia* species and the resulting inflammatory host response have been implicated in both of these disease states. The nature of genomic rearrangements in each species may influence its ability to adapt and cause disease in a specific host niche. Thus, studying the mechanisms of karyotype evolution is an important step towards understanding the evolution of the *Malassezia* species complex.

Kinetochore proteins serve as useful tools in the identification of centromeres due to their centromere-exclusive localization. CENP-A replaces histone H3 in the centromeric nucleosomes. This has been shown as a reduction in histone H3 levels at the centromeres in *C. lusitaniae* (Kapoor et al., 2015) as well as in a human neocentromere (Lo et al., 2001). These CENP-A nucleosomes act as a foundation to recruit CENP-C, the KMN (KNL1C-MIS12C-NDC80C) network, and other kinetochore proteins (Musacchio and Desai, 2017).

In this study, we experimentally validated all of the eight centromeres of *M. sympodialis* using the Mtw1 protein (Mis12 in *S. pombe*), a subunit of the KMN complex, as the kinetochore marker. The Mis12 complex proteins are evolutionarily conserved outer kinetochore proteins that link the chromatin-associated inner kinetochore proteins to the microtubule-associated outer kinetochore proteins. Members of the Mis12 complex localize to centromeres in other organisms (Goshima et al., 1999, Goshima et al., 2003, Westermann et al., 2003, Roy et al., 2011). Recent studies suggest that the protein domains associated with the Mis12 complex members are exclusive to kinetochore proteins and are not detected in any other proteins making them attractive tools to identify centromere sequences (Tromer et al., 2019). Using the features of centromeres of *M. sympodialis* and newly generated chromosome-level genome assemblies, we predicted centromeres in related *Malassezia* species carrying seven, eight, or nine chromosomes and experimentally validated the centromere identity in representative species of each karyotype, belonging to different *Malassezia* clades. We employed gene synteny conservation across these centromeres to understand the transition between these states from an inferred ancestral state of 9 chromosomes. Based on our results, we propose that centromere loss by two distinct mechanisms drives karyotype diversity.

## Results

### Chromosome number varies in the *Malassezia* species complex

Previous reports based on pulsed-field gel electrophoresis (PFGE) have suggested that chromosome number varies within the *Malassezia* species complex. The early diverged species *M. slooffiae* of Clade C was reported to have 9 chromosomes (Boekhout et al., 1998). Clade B *Malassezia* species are reported to have 9 (*M. globosa* and *M. restricta)*, 8 (*M. sympodialis*), or 6 chromosomes (*M. pachydermatis*). Among the Clade A species, *M. obtusa* and *M. furfur* CBS14141 were reported to have 7 chromosomes each (Boekhout and Bosboom, 1994, Zhu et al., 2017). A high-quality reference genome is a prerequisite to understanding the rearrangements associated with chromosome number variation. Additionally, this will also assist in resolving ambiguities in PFGE-based estimates of chromosome number when similar-sized chromosomes are present. Complete genome assemblies were not available for many of the species with reported numbers of chromosomes. To obtain better-assembled reference genomes, we sequenced the genomes of *M. slooffiae* and *M. globosa* as representatives of the 9-chromosome state, and *M. furfur* as a representative of the 7-chromosome state using the PacBio SMRT sequencing technology (Figure 1, Figure 1-figure supplement 1).

**Figure 1.**
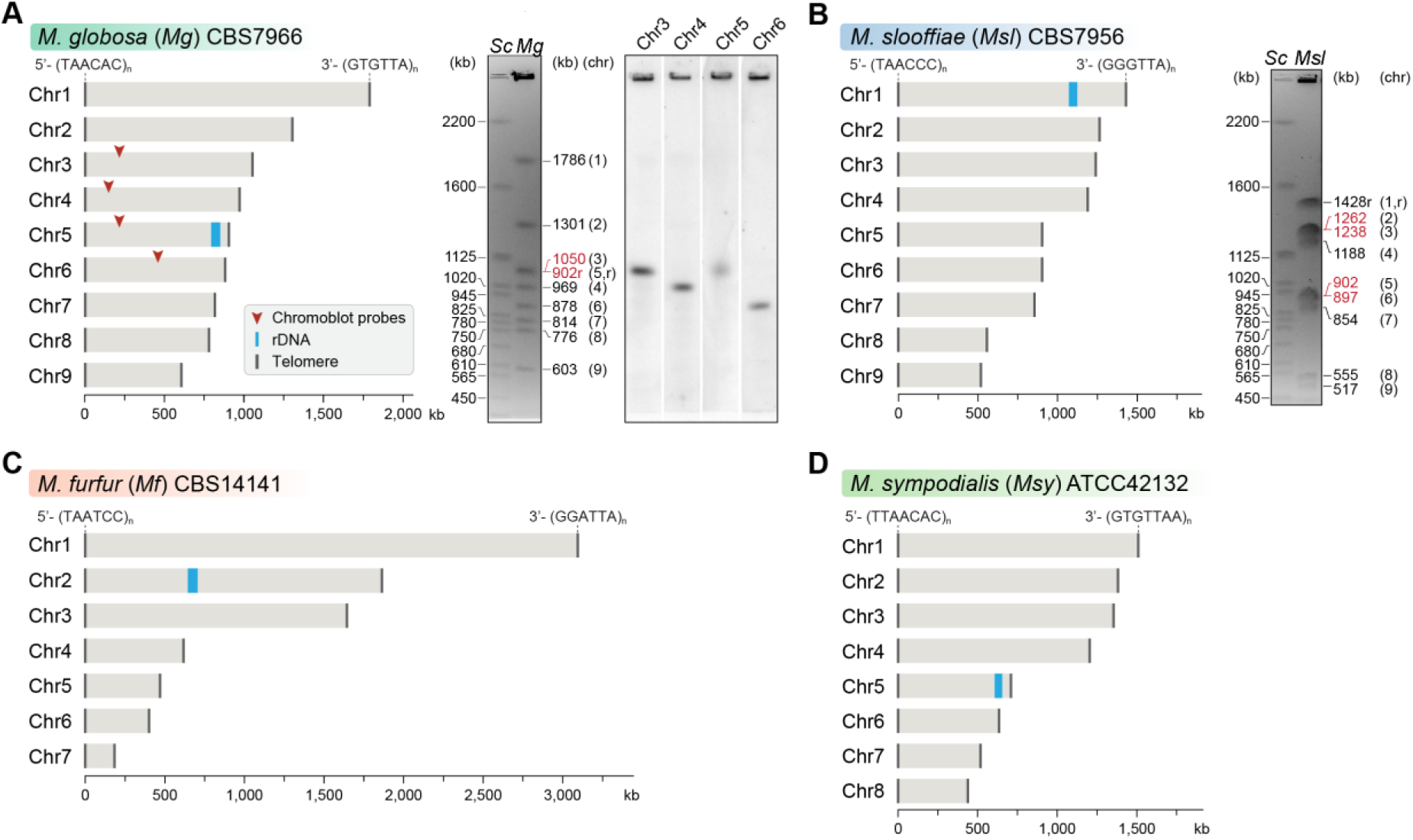
Genome assembly and karyotype diversity in representative *Malassezia* species. The genomes of *M. globosa* (A), *M. slooffiae* (B), and *M. furfur* (C) were sequenced and assembled in this study, while the genome assembly of *M. sympodialis* (D) was reported earlier (Zhu et. al., 2015) and is shown for comparison. In each panel, bar plots represent the assembled chromosomes of the indicated *Malassezia* species, with the telomeres and the ribosomal DNA (rDNA) marked as dark grey and blue bars, respectively. Telomere-repeat motifs are shown at the 5’- and 3’-ends of the Chr1 in each species. Electrophoretic karyotypes of *M. globosa* (*Mg*) and *M. sloffiiae* (*Msl*), are shown in (A) and (B) respectively, with chromosome sizes estimated from the genome assembly. Chromosomes of *Saccharomyces cerevisiae* (*Sc*) served as size markers. The chromosome containing the rDNA (marked with an “r”), in *M. globosa,* co-migrates with Chr3. This was assessed by chromoblot hybridization using unique sequences from Chr3, Chr4, Chr5, and Chr6 as probes (regions indicated by red arrowheads). Chromosomes of similar size (denoted in red) migrate together in the gel and appear as a doublet band (i.e. MgChr3-MgChr5, MslChr2-MslChr3, and MslChr5-MslChr6).

The *M. globosa* genome was completely assembled into 9 contigs with telomeres on both ends (BioSample accession SAMN10720087). We validated these by assigning each band on the pulsed-field gel with the sizes from the genome assembly and further confirmed these by chromoblot analysis following PFGE. This analysis shows that chromosome 5 contains the rDNA locus and migrates higher than the expected size of 902 kb, as a diffuse ensemble of different sizes along with chromosome 3 (Figure 1A). The assembled genome of *M. slooffiae* has 14 contigs of which 9 contigs have telomeres on both ends, indicative of 9 chromosomes (BioSample accession SAMN10720088). Each of the 9 contigs could be assigned to the bands observed in the pulsed-field gel (Figure 1B). For *M. furfur*, the final genome assembly consisted of 7 contigs with telomeres on both ends and matched the expected chromosome sizes obtained from an earlier PFGE analysis of CBS14141 (Figure 1C). The complete genome assembly of *M. sympodialis* reported earlier is distributed into 8 chromosomes with telomere repeats on both ends (Figure 1D), and serves as a representative of an 8-chromosome state in this study.

Changes in chromosome number are always associated with birth or loss of centromeres to stabilize the karyotype in organisms with monocentric chromosomes. To understand the transitions between these different karyotypic states observed in the *Malassezia* species complex, we sought to experimentally validate centromeres in species representative of each karyotype.

### Kinetochores cluster and localize to the nuclear periphery in *M. sympodialis*

Organisms with point centromeres possess Ndc10, Cep3, and Ctf13 of the CBF3 complex, a cognate protein complex specific to point centromeres. None of these point centromere-specific proteins could be detected in *M. sympodialis*. However, we could detect homologs of CENP-A, CENP-C, and most of the outer kinetochore proteins in the genome of *M. sympodialis* (Figure 2A and Figure 2-figure supplement 1). We functionally expressed an N-terminally GFP-tagged Mtw1 protein (Protein ID: SHO76526.1) from its native promoter and the expression of the fusion protein was confirmed by western blotting (Figure 2B). Upon staining with anti-GFP antibodies and DAPI (4′,6-diamidino-2-phenylindole), we could detect punctate localization of Mtw1 at the nuclear periphery (Figure 2C) consistent with the clustered kinetochore localization observed in other yeasts (Goshima et al., 1999, Euskirchen, 2002, Roy et al., 2011). Live-cell images of MSY001 (GFP-*MTW1*) cells revealed that the kinetochores (GFP-Mtw1) remained clustered throughout the cell cycle, starting from unbudded G1 cells in interphase to large budded cells in mitosis (Figure 2D).

**Figure 2.**
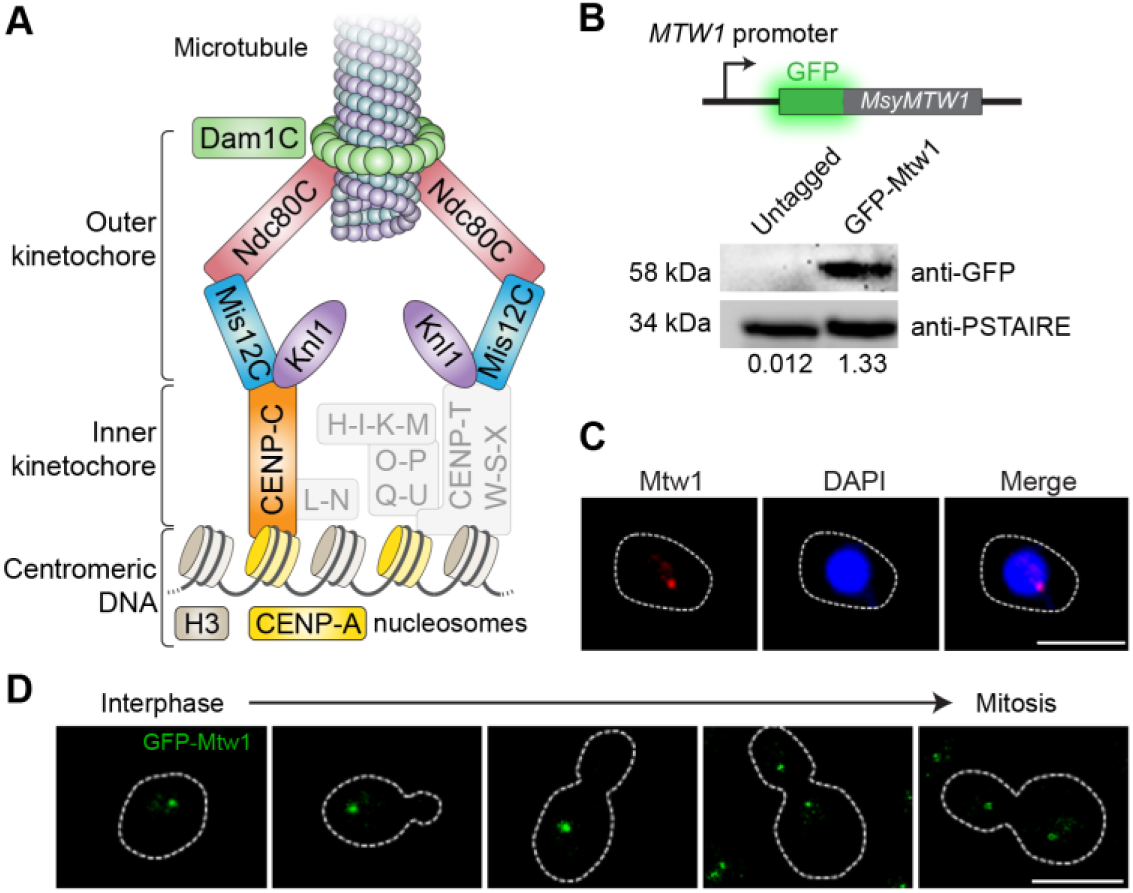
Kinetochores cluster and localize at the nuclear periphery in *M. sympodialis*. (A) Schematic of the kinetochore organization of *M. sympodialis*. Gray boxes indicate proteins absent in *M. sympodialis*. The outer kinetochore protein Mtw1 (a component of Mis12C) served as the kinetochore marker in the present study. (B) Line diagram representation of Mtw1 tagged with GFP at the N-terminus. Immunoblot analysis of whole cell lysates prepared from the untagged *M. sympodialis* strain (ATCC42132) and GFP-Mtw1 expressing cells (MSY001) probed with anti-GFP antibodies and anti-PSTAIRE antibodies. PSTAIRE served as a loading control. Relative intensity values normalized to PSTAIRE are indicated below each lane. (C) Logarithmically grown MSY001 cells expressing GFP-Mtw1 were fixed and stained with DAPI (blue) and anti-GFP antibodies (pseudo-colored in red). Scale bar, 2.5 µm. (D) Cell cycle stage-specific localization dynamics of GFP-Mtw1. Scale bar, 2.5 µm.

### Mtw1 is localized to a single region at the GC minima of each *M. sympodialis* chromosome

Having identified Mtw1 as an authentic kinetochore protein, we performed ChIP-sequencing using the GFP-Mtw1 expressing strain of *M. sympodialis* (MSY001). Mapping the reads to the reference genome of *M. sympodialis* strain ATCC42132 (Zhu et al., 2017) identified one significantly enriched locus on each of the eight chromosomes (Figure 3A). The length of the Mtw1-enriched centromere regions identified from the ChIP-seq analysis ranges between 3167 bp and 5143 bp with an average length of 4165 bp (Table S1). However, the region of maximum Mtw1 enrichment on each chromosome (based on the number of sequenced reads aligned) mapped to the intergenic region harboring the GC trough (approximately 1 kb long), which was previously predicted to be the centromeres of *M. sympodialis* (Figure 3B) (Zhu et al., 2017). The regions of Mtw1 enrichment span beyond the core centromeres and include active genes located proximal to these troughs (Figure 3B, Figure 3-figure supplement 1A). However, these ORFs do not show consensus features such as the orientation of transcription or functional classification. We validated this enrichment by ChIP-qPCR analysis with primers homologous to the predicted centromeres compared to a control region distant from the centromere (Figure 3-figure supplement 1B).

**Figure 3.**
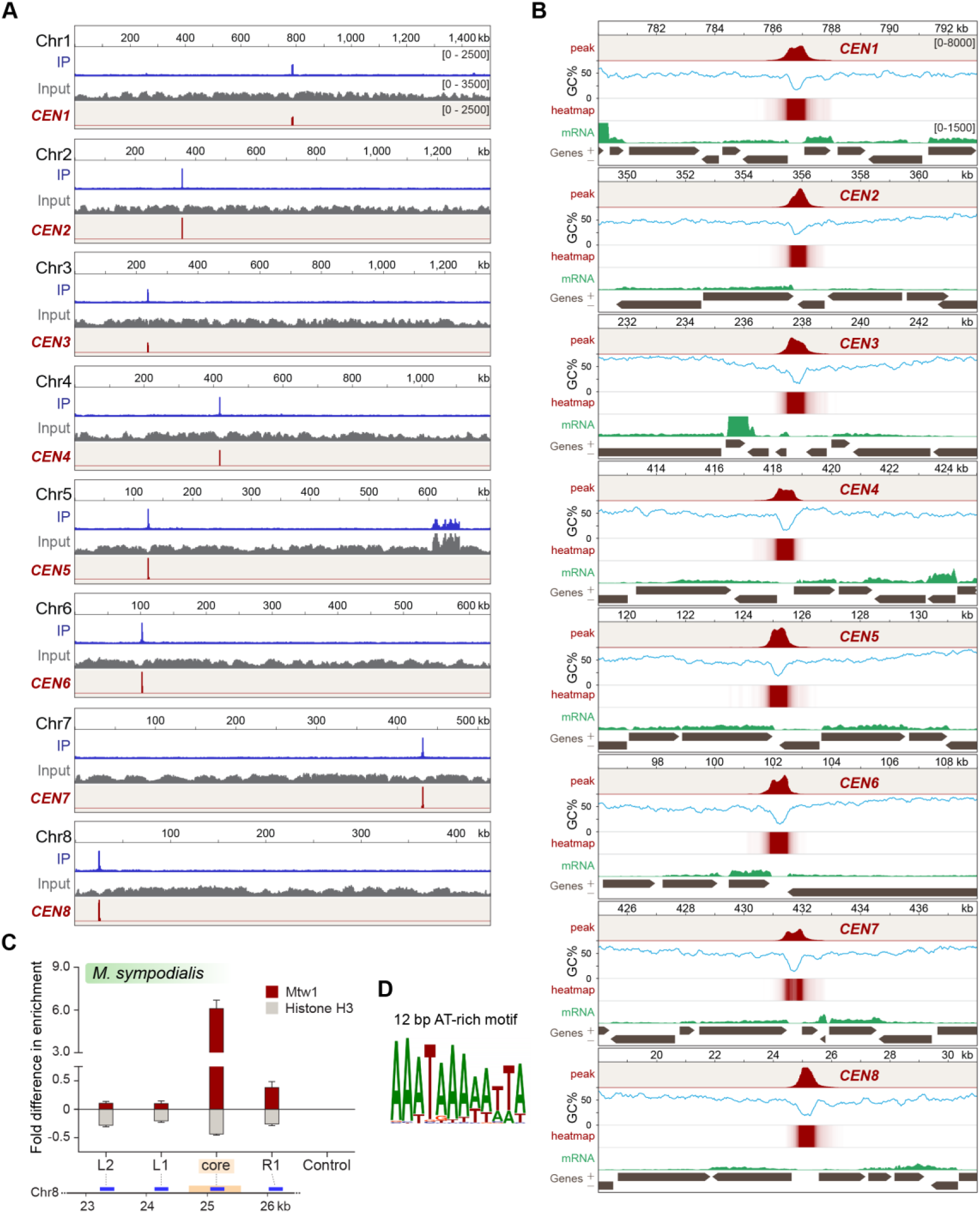
Localization of Mtw1to single-peaks identifies centromeres on each of the eight chromosomes of *M. sympodialis.* (A) GFP-Mtw1 ChIP-seq reads were mapped to each chromosome. The *x*-axis indicates chromosomal coordinates (in kb) and the *y*-axis indicates read depth. “Input”, reads from total DNA; “IP,” reads from immunoprecipitated sample; *CEN,* Mtw1-enriched regions derived by subtracting input reads from those of the IP sample (peak values 0-2500). Additional peaks observed in both IP and input tracks on Chr5 are from the rDNA locus. (B) A 13 kb-window of Mtw1 enrichment profile (*CEN*, represented as peaks and heat-map in two different tracks, red) plotted along with the GC content (%GC, blue) and regions of transcription (RNA-seq, green). Numbers mentioned in the topmost track in every panel indicate chromosomal coordinates (in kb). The scales in the *y*-axis are as follows: *CEN* (0-8000), %GC (0-75), RNA-seq reads (0-1500). Gray arrows in each panel indicate predicted ORFs based on RNA-seq data with arrowheads pointing the direction of transcription of the corresponding gene, also marked as ‘+’ and ‘-’ in the axis label. (C) Fold difference in Mtw1 and histone H3 enrichment at *CEN8* as compared to a non-centromeric control region (190 kb away on the right of *CEN1*) by qPCR analysis. Schematic of a 4 kb region of Chr8 with *CEN8* core (yellow box) is depicted below the graph. Blue lines indicate regions assayed by PCR: core-region corresponding to the GC trough, L1 and R1-750 bp away from the core, L2-1500 bp away from the core, and non-centromeric control region (190 kb away from centromere in Chr1). The *x*-axis indicates regions across the *CEN8* probed by PCR and *y*-axis indicates fold difference in the enrichment of Mtw1 and histone H3 as compared to the control region. Error bars indicate standard deviation (SD). (D) Logo of the consensus DNA sequence identified from *M. sympodialis* centromeres, graphically represented with the size of the base corresponding with the frequency of occurrence.

### Histone H3 is depleted at the core centromere with active genes at the pericentric regions in *M. sympodialis*

The presence of CENP-A nucleosomes at the kinetochore should result in reduced histone H3 enrichment at the centromeres when compared to a non-centromeric locus. To test this, we performed ChIP with anti-histone H3 antibodies and analyzed the immunoprecipitated (IP) DNA by qPCR. As compared to a control ORF region unlinked to the centromere (190 kb away from *CEN1*), the pericentric regions flanking the core centromere showed a marginal reduction in histone H3 enrichment that was further reduced at the core that maps to the GC trough with the highest enrichment of the kinetochore protein Mtw1. That the core centromere region showing the maximum depletion of histone H3 coincided with the regions most enriched with Mtw1 further supports that histone H3, in these regions, is possibly replaced by its centromere-specific variant CENP-A (Figure 3C).

### The short regional centromeres of *M. sympodialis* are enriched with a 12 bp long AT-rich consensus sequence motif

To understand the features of *M. sympodialis* centromeres, we analyzed the centromere DNA sequences for the presence of consensus motifs or structures such as inverted repeats. PhyloGibbs-MP (Siddharthan et al., 2005, Siddharthan, 2008) predicted a 12 bp long AT-rich motif common to all of the centromere sequences of *M. sympodialis* (Figure 3D). We swept the Position Weight Matrix (PWM) from the PhyloGibbs-MP output across each chromosome of *M. sympodialis* and counted the number of motif predictions in sliding 500 bp window, sliding by 100 bp at a time. Sites with log-likelihood-ratio (LLR) of >7.5 were counted as motif predictions. The LLR is the natural logarithm of the ratio of the likelihood of the 12 bp substring arising as a sample from the PWM, to the likelihood of it being generic “background”. In each case, the global peak coincides with the centromere (Figure 3-figure supplement 2A). In each chromosome, the centromere region shows between 7 and 13 motif matches, while no other 500 bp window shows more than 3 matches. This suggests that the AT-rich motif is more enriched at the centromeres than at any other region in the *M. sympodialis* genome (Figure 3-figure supplement 2A). To ensure that this is not an artifact of the GC-poor nature of the centromere, we repeated the analysis with a synthetic shuffled PWM, created by scrambling the order of the columns of the original PWM (that is, scrambling the positions in the motif while keeping the corresponding weight vectors the same). This shuffled motif showed more matches in the centromeres than what is seen in the non-centromeric genomic sequence, but significantly fewer than the authentic centromeric sequences in most chromosomes (Figure 3-figure supplement 2B). A dot-plot analysis was performed to detect the presence of any direct or inverted-repeat structure associated with the centromeres in *M. sympodialis*. Analysis of all of the centromere sequences along with 5 kb flanking sequences using SyMap confirmed the lack of direct/inverted repeat structures (Figure 3-figure supplement 2C).

In the absence of any centromere exclusive DNA sequence, the unique and distinguishing features of centromere regions in *M. sympodialis* are an AT-rich core region of <1 kb (average AT content of 78% as compared to the genome average of 41.5%) enriched with the 12 bp motif (Figure 3-figure supplement 3) in a kinetochore protein-bound region of 3 to 5 kb. As expected, the kinetochore bound region contains a reduced level of histone H3.

### Centromeres in *M. furfur, M. slooffiae,* and *M. globosa* map to chromosomal GC minima

Using the unique centromere features identified in *M. sympodialis*, we predicted one centromere locus on each of the 7 *M. furfur* chromosomes and these all map to chromosomal GC troughs (Figure 4-figure supplement 1A, Table S2). Similar to *M. furfur*, we predicted the centromeres in *M. slooffiae, M. globosa,* and *M. restricta*, each of which contains 9 chromosomes (Figure 4-figure supplement 1B-D, Table S2). Each of the predicted centromeres showed enrichment of the 12 bp AT-rich motif identified with *M. sympodialis* centromere sequences as compared to other regions in the genomes (Figure 4-figure supplement 2).

To experimentally validate the centromeric loci in *M. furfur*, we functionally expressed the centromeric histone H3 variant CENP-A with a 3xFLAG tag at the C-terminus (Figure 4A). We performed ChIP in strain MF001 (CENP-A-3xFLAG) and analyzed immunoprecipitated DNA by qPCR using primers specific to each of the seven predicted centromeres and a centromere-unlinked control locus 1.3 Mb away from *CEN1*. Enrichment of CENP-A at all seven centromeres over the control locus confirmed that the regions predicted are indeed centromeres in *M. furfur* CBS14141 (Figure 4B, C).

**Figure 4.**
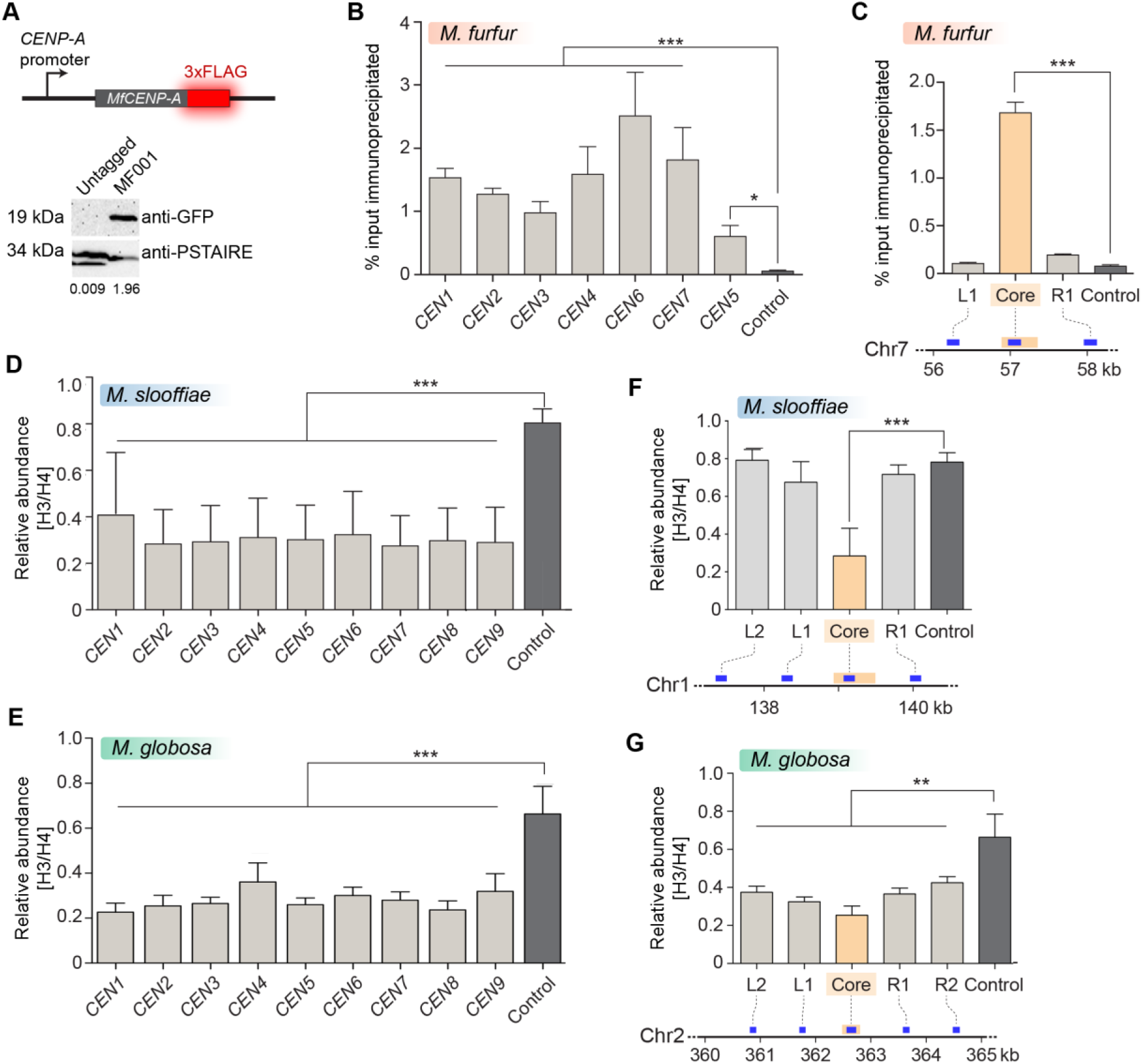
Centromeres in *M. furfur*, *M. slooffiae,* and *M. globosa* map to global GC troughs on each chromosome. (A) Schematic of epitope tagging of CENP-A with 3xFLAG at the C-terminus is shown. (Bottom) Immunoblot analysis using whole cell lysates prepared from the untagged wild-type *M. furfur* (CBS14141) cells and CENP-A-3xFLAG expressing cells (MF001) probed with anti-FLAG antibodies and anti-PSTAIRE antibodies. PSTAIRE served as the loading control. (B) Abundance of CENP-A at each of the predicted *M. furfur* centromeres by qPCR analysis of DNA immunoprecipitated with anti-FLAG affinity gel in MF001 cells expressing CENP-A-3xFLAG. The *x*-axis indicates individual *CEN* regions assayed with primers homologous to the GC troughs on each chromosome that were predicted as centromeres (see Table S7 for primer sequences). The non-centromeric control maps to a region 1.3 Mb away from predicted *CEN1*. The *y*-axis indicates enrichment of CENP-A estimated as the percentage of input immunoprecipitated. (C) Abundance of CENP-A across *MfCEN7* by ChIP-qPCR analysis in MF001 cells. Schematic representation of a 2 kb region is shown below the graph. Yellow bar indicates the centromere core of *CEN7* corresponding to the GC trough. Blue bars indicate regions analyzed by qPCR: L1 and R1-750 bp away from the centromere core. ‘Control’ refers to a region 1.3 Mb away from *CEN1*. The CENP-A enrichment is plotted along the *y*-axis as the percentage of input immunoprecipitated. (D, E) Comparison of relative abundance of histone H3 compared to histone H4 at the predicted centromeres to a non-centromeric control locus in *M. slooffiae* and *M. globosa* respectively. Enrichment was estimated as the percentage of input immunoprecipitated with histone H3 and histone H4 antibodies and their ratio, as indicated in *y-*axis, is plotted as relative enrichment. The *x-*axis indicates centromeres in each species and these were assayed with primers homologous to GC troughs in each chromosome that were predicted as the centromeres (see Table S7 for primer sequences). The control region unlinked to the centromere corresponds to a locus 630 kb away from *CEN1* in *M. slooffiae* and 416 kb away from *CEN2* in *M. globosa* respectively. (F) The relative abundance of histone H3 compared to histone H4 across *MslCEN1* by qPCR analysis of the DNA immunoprecipitated using histone H3 and histone H4 antibodies. Schematic of *MslCEN1* locus is shown below the graph. Yellow bar indicates the *CEN1* core region corresponding to the GC trough. Blue bars indicate regions analyzed by qPCR: L1 and L2 map to regions 750 bp and 1.5 kb to the left of the *CEN1* core; R1 maps to a region 750 bp to the right of the *CEN1* core. The control region corresponds to a locus 630 kb away from the *CEN1* core. The ratio of enrichment of histone H3 to that of histone H4 is plotted as the relative enrichment in *y*-axis. (G) The relative abundance of histone H3 compared to histone H4 across *MgCEN2* by qPCR analysis of the DNA immunoprecipitated using histone H3 and histone H4 antibodies. Schematic of a 5 kb region containing *MgCEN2* is shown below the graph. Yellow bar indicates the *CEN2* core region corresponding to the GC trough. Blue bars indicate regions analyzed by qPCR: L1 and L2 indicate regions 750 bp and 1.5 kb to the left of the *CEN2* core; R1 and R2 indicate regions 750 bp and 1.5 kb to the right of the *CEN2* core. The control region corresponds to a locus 416 kb away from the *CEN2* core. The ratio of enrichment of histone H3 to that of histone H4 is plotted as the relative enrichment in the *y*-axis. Error bars in figures B-G indicate standard deviation (SD); Statistical significance was tested by one-way ANOVA: * significant at P < 0.05, *** significant at P < 0.001.

Given the lack of genetic manipulation methods for *M. slooffiae* and *M. globosa*, we tested the enrichment of histone H3 at the predicted centromeres in these two species. All of the nine centromeric loci in these two species showed a reduced histone H3 level when compared to a control locus unlinked to centromeres (Figure 4D, E). Furthermore, upon analyzing the enrichment profile at one centromere (*CEN1*) in *M. slooffiae*, we observed a reduction in the enrichment levels of histone H3 at the GC troughs as compared to the flanking regions (Figure 4F). In the case of *M. globosa*, the regions spanning a centromere (*CEN2*) also depicted a similar reduction in the histone H3 levels (Figure 4G). Taken together, the significant reduction in the histone H3 levels at the predicted centromeres, indicative of the presence of CENP-A, suggests that these putative centromere regions are indeed *bona fide* centromeres in these species.

### Centromere loss by breakage resulted in chromosome number reduction in *M. sympodialis*

Synteny of genes across centromeres is largely conserved in closely related species (Byrne and Wolfe, 2005, Padmanabhan et al., 2008, Yadav et al., 2018b). To understand the transition between different chromosome number states, we analyzed the conservation of gene synteny across centromeres in these species. By mapping gene synteny at the centromeres of *M. globosa* and *M. slooffiae* (each carries 9 chromosomes), compared with *M. sympodialis* (containing 8 chromosomes), we found complete gene synteny conservation in 8 of the 9 centromeres (Figure 5A, B). Thus, syntenic regions of all 8 *M. sympodialis* centromeres are present in the genomes of *M. globosa* and *M. slooffiae*. In the case of *M. restricta*, 7 putative centromeres are completely syntenic with *M. sympodialis* centromeres and one centromere retained partial gene synteny (Table S3). However, no gene synteny conservation was observed at the centromeres of Chr2 in *M. globosa,* Chr5 in *M. slooffiae,* or Chr8 in *M. restricta* (Table S3), indicative of loss of a centromere during the transition from the 9-chromosome state to the 8-chromosome state.

**Figure 5.**
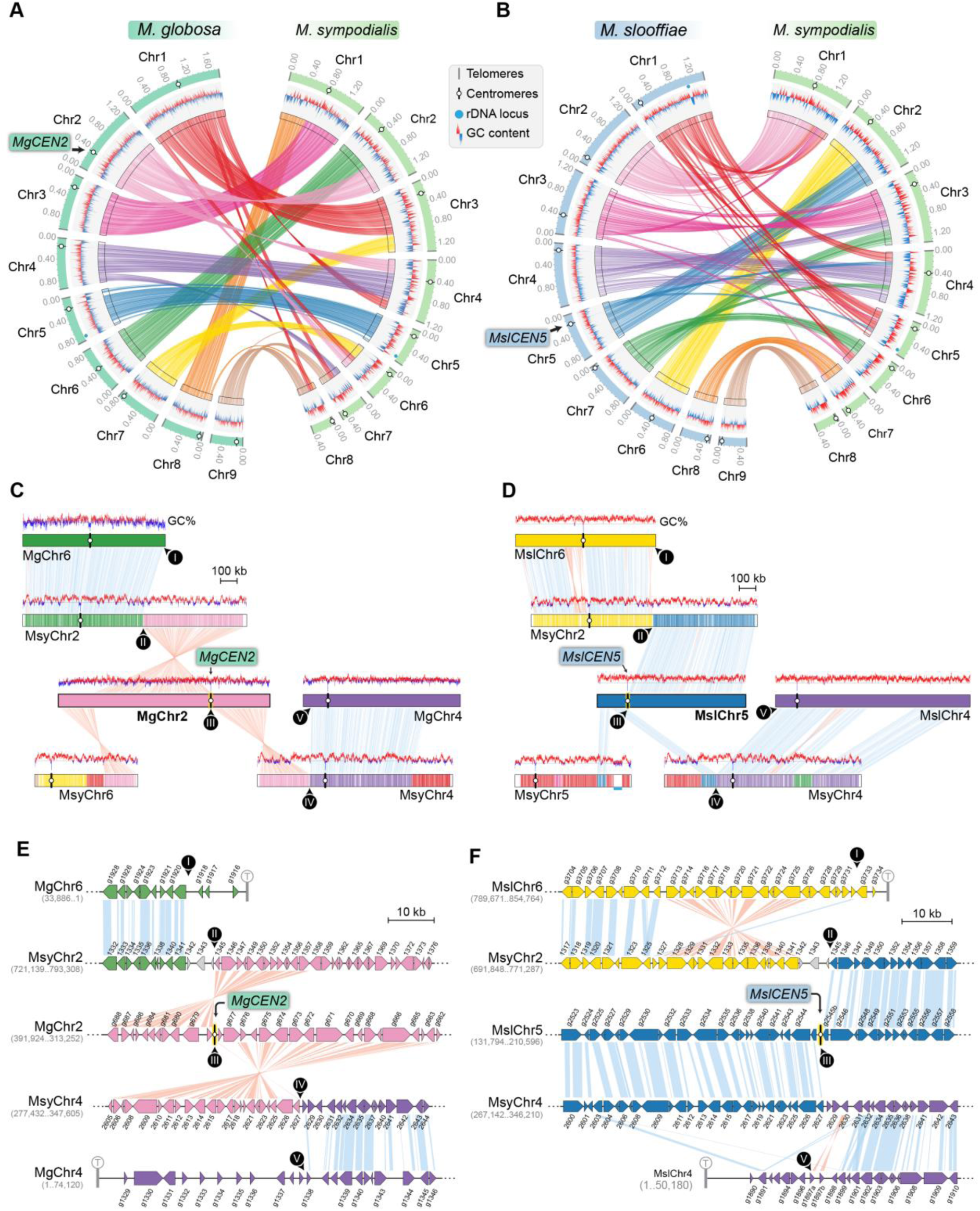
*MgCEN2* and *MslCEN5* map to a synteny breakpoint in *M. sympodialis*. (A, B) Circos plots depicting the conserved gene synteny blocks between *M. globosa* and *M. sympodialis* chromosomes and *M. slooffiae* and *M. sympodialis* chromosomes. Tracks from outside to inside represent positions of centromeres and telomeres, GC content (plotted as blue and red lines indicating GC content below or above genome average, calculated in 0.4 kb non-overlapping windows), and colored connectors indicate regions of conserved gene synteny between the two species. (C, D) Linear chromosome plots depicting syntenic regions between Chr2 of *M. globosa* and Chr5 of *M. slooffiae* respectively with chromosomes of *M. sympodialis.* GC content (in %) is shown as red/blue lines above each chromosome. Labels in black circles mark the gene synteny breakpoints. Synteny breakpoint at *MgCEN2* and *MslCEN5* is marked as III. The regions on MsyChr2 and MsyChr4 where the homologs of ORFs flanking the breakpoint are located, are marked II and IV respectively. Labels I and V indicate gene synteny conservation on the other side of the fusion points II and IV on MsyChr2 and MsyChr4 as compared to *M. globosa* and *M. slooffiae* chromosomes respectively. (E, F) Zoomed-in image of the gene synteny breakpoint at *MgCEN2* and *MslCEN5* representing the conservation of genes flanking these centromeres in *M. sympodialis* chromosomes at the ORF level.

The GC trough corresponding to *MgCEN2/MslCEN5* is flanked by genes that map to MsyChr2 on one arm and MsyChr4 on the other (Figure 5C, D). The centromere region in each of MgChr2 and MslChr5 marks a synteny breakpoint showing no homologous region in the *M. sympodialis* genome indicative of a loss of this centromere DNA sequence. We also observed that the genes flanking the breakpoint are conserved in *M. sympodialis*, suggesting that the specific intergenic region was involved (Figure 5E, F). Evidence for internalization of telomere adjacent ORFs or the presence of interstitial telomere repeats indicative of telomere-telomere fusions was not detected in the *M. sympodialis* genome. These observations strongly support our hypothesis that breakage of *MgCEN2/MslCEN5* (or the orthologous ancestral *CEN*) and fusion of the two acentric arms to other chromosomes resulted in the chromosome number reduction observed between these species.

### Centromere inactivation by sequence divergence and loss of AT-richness resulted in chromosome number reduction in *M. furfur*

To understand the basis of the change in chromosome number from 9 to 7 in *Malassezia* species, we compared the synteny of ORFs flanking the *M. slooffiae* or *M. globosa* centromeres with that of *M. furfur*. Of the 9 centromeres in *M. slooffiae*, three centromeres belonged to conserved gene synteny blocks and four other centromeres showed partial gene synteny conservation in *M. furfur* (Figure 6A, Table S3). A similar pattern of gene synteny conservation was observed between *M. globosa* and *M. furfur* (Figure 6B, Table S3). The genes flanking the remaining two centromeres (*CEN8* and *CEN9*) in *M. slooffiae* were present in conserved gene synteny blocks in the two arms of MfChr3 (Figure 6C). However, the regions corresponding to *CEN8* and *CEN9* in *M. slooffiae* appear to have evolved to decreased AT-richness in *M. furfur.* A similar centromere inactivation mechanism was observed when *CEN8* and *CEN9* of *M. globosa* were compared to the corresponding syntenic regions in *M. furfur* (Figure 6D). These results are suggestive of centromere inactivation by changes in the centromeric DNA sequence in this species (Figure 6-figure supplement 1).

**Figure 6.**
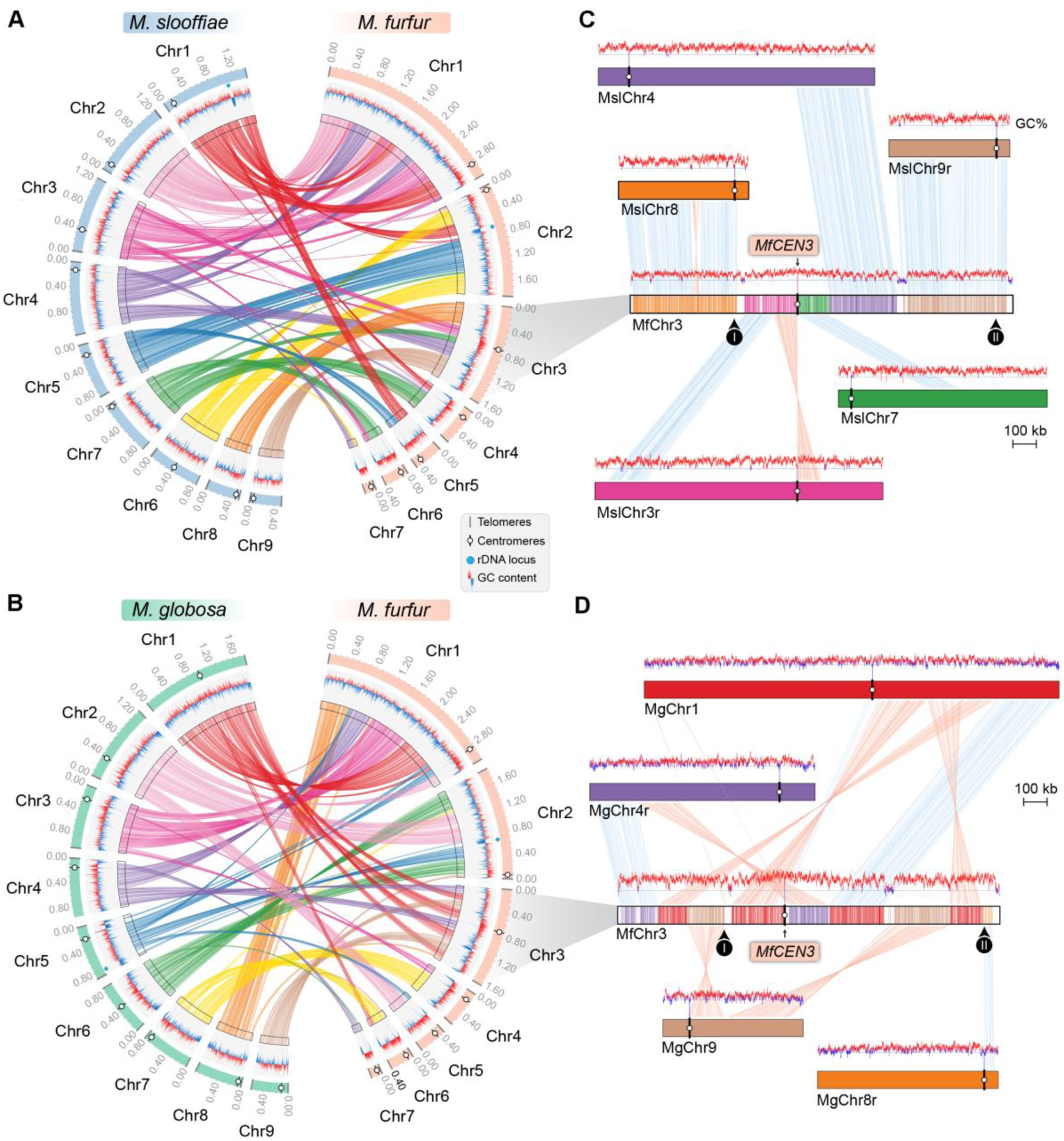
Inactivation of *CEN8* and *CEN9* of *M. slooffiae* and *M. globosa* in MfChr3 resulted in chromosome number reduction in *M. furfur.* (A, B) Circos plots depicting the conserved gene synteny blocks between the *M. slooffiae* and *M. globosa* chromosomes respectively with *M. furfur* chromosomes. Tracks from outside to inside represent positions of centromeres and telomeres, GC content (plotted as blue and red lines indicating GC content below or above genome average, calculated in 0.4 kb non-overlapping windows), and colored connectors indicate regions of conserved gene synteny between the two species. (C) Linear chromosome plot depicting the syntenic regions between chromosome 3 of *M. furfur* and chromosomes 3, 4, 7, 8, and 9 of *M. slooffiae.* GC content (in %) is shown as red/blue lines above each chromosome. Regions corresponding to *MslCEN8* and *MslCEN9* in MfChr3 are marked I and II respectively. (D) Linear chromosome plot depicting the gene synteny conservation between chromosome 3 of *M. furfur* and chromosomes 1, 4, 8, and 9 of *M. globosa*. Regions corresponding to *MgCEN9* and *MgCEN8* in MfChr3 are marked I and II respectively.

### The common ancestral *Malassezia* species contained 9 chromosomes

To trace the ancestral karyotype in *Malassezia*, we predicted centromeres and inferred chromosome numbers in other species of clades A and B based on GC troughs and gene synteny. We identified putative centromeres in *M. dermatis* and *M. nana* in Clade B because of their relatively better-assembled genomes distributed in 18 and 13 scaffolds respectively. Of these, we could predict 8 centromeric regions marked by GC troughs that were also enriched with the 12 bp motif in each species (Figure 7-figure supplement 1A-B, Figure 7-figure supplement 2A-B and Table S2). Furthermore, in both of these species, the 8 putative centromeres shared complete gene synteny conservation with the regions spanning *M. sympodialis* centromeres, indicating that their common ancestor had 8 chromosomes (green circle in Figure 7A, Figure 7-figure supplement 3 and Table S3). To map the common ancestor in Clade B *Malassezia* species, we analyzed regions flanking centromeres of Chr2 of *M. globosa* and Chr8 of *M. restricta,* both of which mapped to the gene synteny breakpoint of the genome of the *Malassezia* species with 8 chromosomes suggesting their common ancestor, named as Ancestor B (Anc. B), also had 9 chromosomes (Figure 7A, Figure 7-figure supplement 3). Based on our centromere predictions in Clade B species and synteny analysis, we propose that centromere breakage would have occurred in the common ancestor of *M. sympodialis*, *M. nana,* and *M. dermatis* after divergence from the common ancestor of *M. globosa* and *M. restricta* which retained a 9-chromosome configuration (Figure 7A, B).

**Figure 7.**
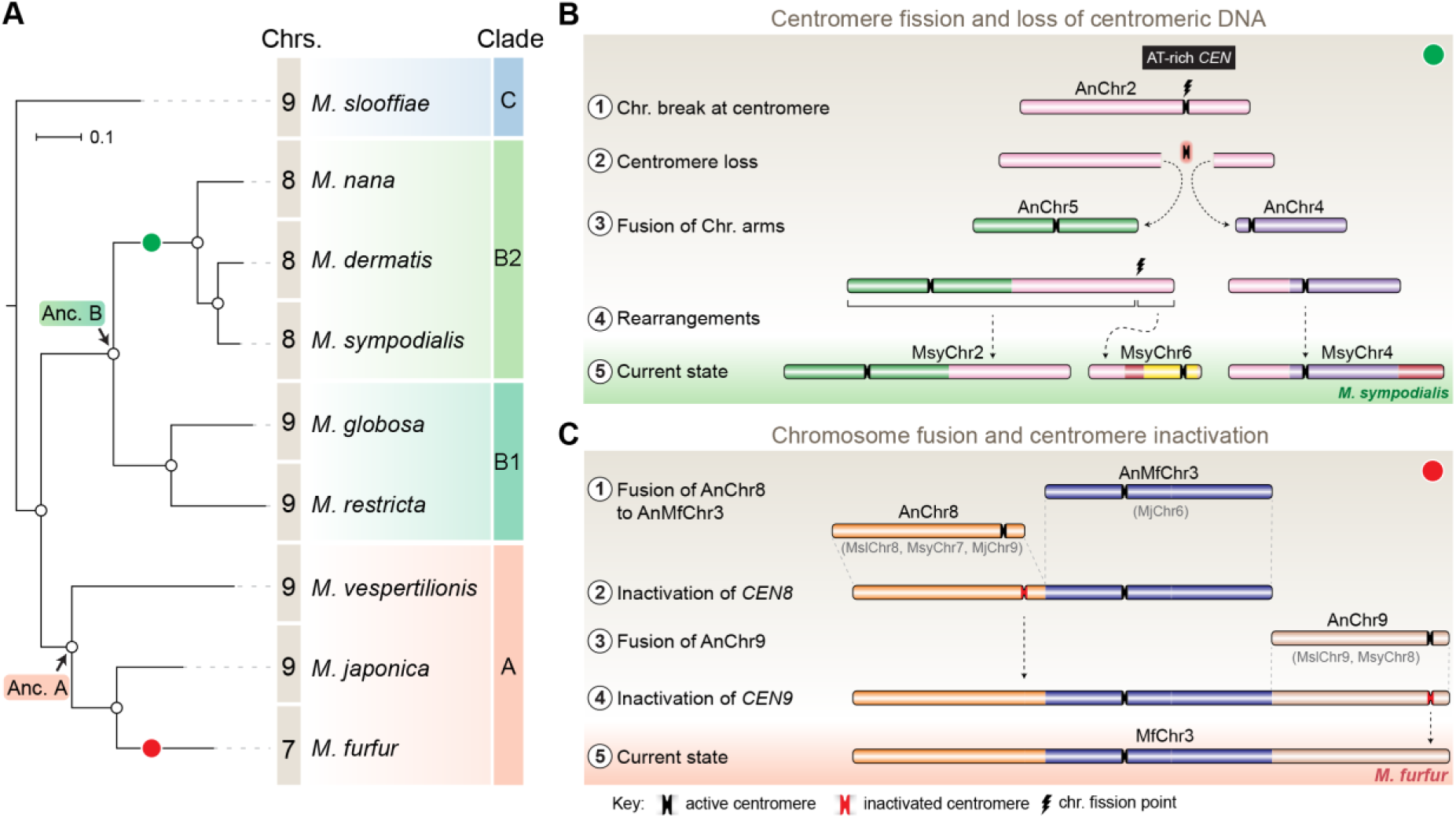
Karyotype evolution by loss of centromere function in *Malassezia* species. (A) Phylogenetic relationships between the *Malassezia* species analyzed in this study are represented and their chromosome numbers shown. Species representing each clade are color coded based on previous reports (Theelen et al., 2018). The chromosome numbers for *M. slooffiae* and *M. globosa* are based on results from this study. In the case of *M. sympodialis, M. restricta,* and *M. furfur*, the chromosome numbers are based on previous reports (Zhu et al., 2017, Senczek et al., 1999, Boekhout and Bosboom, 1994). For *M. dermatis, M. nana*, *M. vespertilionis,* and *M. japonica,* the number of chromosomes were estimated from the predicted number of centromeres. The node corresponding to the ancestral state of Clade A and Clade B are labelled ‘Anc. A’ and ‘Anc. B’ respectively. Green and red circles indicate the origins of karyotypes with 8 and 7 chromosomes respectively from an ancestral state of 9 chromosomes. (B) Schematic of the centromere loss by breakage and the resulting reduction in chromosome number as observed in *M. sympodialis* (represented as the current state). A karyotype with 9 chromosomes (as shown for *M. globosa*) is depicted as the ancestral state. (C) Proposed model of centromere inactivation observed in *M. furfur* as a consequence of fusion of AnChr8 and AnChr9 to the AnMfChr3 equivalent, resulting in a 7-chromosome configuration. The fusion product corresponding to extant MfChr3 is represented as the current state.

As mentioned earlier, *M. furfur* and *M. obtusa* of Clade A contain 7 chromosomes each (Boekhout and Bosboom, 1994, Boekhout et al., 1998). To further understand the karyotype variations within this clade, we predicted the chromosome number in *Malassezia vespertilionis* and *Malassezia japonica* as their genomes are relatively well assembled (Sugita et al., 2003, Lorch et al., 2018). In both of these species, we could predict 9 GC troughs indicative of centromeres of 9 chromosomes (Figure 7-figure supplement 1C-D, Table S2). In the case of *M. japonica*, all of the predicted centromeres (except the centromere of scaffold 7) showed enrichment of the 12 bp motif (Figure 7-figure supplement 2C). However, the 12 bp motif was found to be enriched in all of the predicted centromeres of *M. vespertilionis* (Figure 7-figure supplement 2D). The presence of species with 9 chromosomes in Clade A suggests that the ancestral state in this clade, named Ancestor A (Anc. A) also contained 9 chromosomes (Figure 7A).

We identified 9 centromeres in *M. slooffiae*, the only species in Clade C with a well-assembled genome. The presence of species with 9 chromosomes in each of the three clades of *Malassezia* species, the conservation of gene synteny across orthologous centromeres, and the similar centromere features shared by all 9 species analyzed in this study suggest that *Malassezia* species diverged from a common ancestor that had 9 chromosomes with short regional centromeres enriched with the 12 bp AT-rich DNA sequence motif.

## Discussion

In this study, we experimentally validated the chromosome number in *M. slooffiae* and *M. globosa* by PFGE analysis. We sequenced and assembled the genomes of *M. slooffiae*, *M. globosa*, and *M. furfur* and compared each one with the genome of *M. sympodialis* to understand the karyotype differences observed in members of the *Malassezia* species complex. These species represent each of the three major clades of *Malassezia* species with chromosome numbers ranging from 7 to 9. Because centromere loss or gain directly influences the chromosome number of a given species, we experimentally identified the centromeres of these representative species to understand the mechanisms of karyotype diversity. Kinetochore proteins are useful tools in identifying the centromeres of an organism. Localization of the evolutionarily conserved kinetochore protein Mtw1 tagged with GFP suggested that kinetochores are clustered throughout the cell cycle in *M. sympodialis*. ChIP-sequencing analysis identified short regional (< 5 kb long) centromeres in *M. sympodialis* that are depleted of histone H3, and are enriched with an AT-rich sequence motif. Identification of centromeres in *M. slooffiae*, *M. globosa,* and *M. furfur* further suggested that centromere properties are conserved across these *Malassezia* species. By predicting putative centromere in five other species along with four species with experimentally mapped centromeres described above across three clades of *Malassezia*, we concluded that an AT-rich centromere core of < 1 kb in length enriched with the 12 bp sequence motif is a potential signature of centromeres in nine *Malassezia* species analyzed in this study. Comparative genomics analysis revealed two major mechanisms of centromere inactivation resulting in karyotype change. The presence of a 9-chromosome state in each of the three clades along with conserved centromere features and conserved gene synteny across these centromeres, helped us to infer that the ancestral *Malassezia* species had 9 chromosomes with short regional centromeres with an AT-rich core enriched with the 12 bp sequence motif.

Centromeres in the *Malassezia* species complex represent the first example of short regional centromeres in basidiomycetes. Centromeres reported in other basidiomycetes, such as those in *Cryptococcus* and *Ustilago* species, are of the large regional type (Yadav et al., 2018b). *Malassezia* species analyzed in this study have a significantly smaller genome (< 9 Mb) as compared to other basidiomycetes and lack RNAi machinery. The occurrence of short regional centromeres in RNAi-deficient *Malassezia* species is in line with a previous finding wherein a reduction in centromere size was observed in RNAi-deficient basidiomycete species as compared to their RNAi-proficient relatives (Yadav et al., 2018b). With the presence of clustered kinetochores across cell cycle stages, and the absence of key genes encoding the RNAi machinery, these *Malassezia* species resemble ascomycetes such as many of the CTG clade species with short regional centromeres (Sanyal et al., 2004, Nakayashiki et al., 2006, Padmanabhan et al., 2008, Kapoor et al., 2015, Chatterjee et al., 2016, Yadav et al., 2018a), rather than the basidiomycetes with large regional centromeres. By combining these features, we conclude that the genome size and the presence of complete RNAi machinery could be the determinants of the centromere type of a species irrespective of the phylum it belongs to.

Based on the binding pattern of the kinetochore protein across the *M. sympodialis* genome, the 3 to 5 kb long region can be divided into two domains: (a) an AT-rich *CEN* core that maps to the intergenic region containing the GC trough showing maximum kinetochore binding (< 1 kb) and (b) the regions flanking the core that show basal levels of kinetochore protein binding. We observed conservation of the 12 bp AT-rich motif in the centromere core across the nine *Malassezia* species. It should also be noted that the 12 bp motif is significantly enriched at the centromeres but not exclusive to the centromeres as it is detected across the chromosomes. We did not observe any orientation bias of these motifs at the centromeres. The functional significance of the frequent occurrence of this motif at centromeres is unknown. It will be intriguing to test the roles played by this motif, the core, and the flanking sequences in centromere function. Testing these domains for centromere function *in vivo* by making centromeric plasmids in various *Malassezia* species is challenging at present due to technical limitations. Other than *M. sympodialis*, *M. pachydermatis,* and *M. furfur*, no other *Malassezia* species have been successfully transformed (Ianiri et al., 2016, Celis et al., 2017, Ianiri et al., 2017b). Moreover, in all of these cases, the genetic manipulations are performed by *Agrobacterium*-mediated transconjugation, which is not amenable for the introduction of circular plasmids.

The centromeres in *M. sympodialis* contain transcribed ORFs, as documented in centromeres of rice, maize, and *Zymoseptoria tritici* (Nagaki et al., 2004, Wang et al., 2014). In contrast to these cases, our read count analysis did not reveal any significant difference in the transcription (RPKM values) of centromere associated ORFs to that elsewhere in the genome of *M. sympodialis*. We posit that the 12 bp AT-rich motif sequences could facilitate the transcription of these genes by recruiting transcription factors with a possible role in kinetochore assembly. Binding of Cbf1 at CDEI in *S. cerevisiae* centromeres and Ams2 at the central core sequences of *S. pombe* centromeres are classic examples of transcription factors facilitating kinetochore stability in fungal systems (Hemmerich et al., 2000, Chen et al., 2003). A fine regulation of transcription by Cbf1, Ste12, and Htz1 and the resulting low-level cenRNAs have been implicated in proper chromosome segregation in budding yeast (Ohkuni and Kitagawa, 2011, Ling and Yuen, 2019). These studies reinforce the role of transcription in centromere function irrespective of centromere structure.

In this study, we report three high-quality chromosome-level genome assemblies and identified centromeres in nine *Malassezia* species, representing all of the three *Malassezia* clades with differing numbers of chromosomes. This will serve as a rich resource for comparative genomics in the context of niche adaptation and speciation. Analysis of gene synteny conservation across centromeres using these genomes revealed breakage at the centromere as one of the mechanisms resulting in a karyotype change between closely related species - those with 9 chromosomes, such as *M. slooffiae* and *M. globosa,* and those with 8 chromosomes like *M. sympodialis*. Gene synteny breakpoints adjacent to the centromeres have been reported in *C. tropicalis* that has seven chromosomes - one less than *C. albicans* (Chatterjee et al., 2016). Centromere loss by breakage was proposed to have reduced the *Ashbya gossypii* karyotype by one when compared to the pre-whole genome duplication ancestor (Gordon et al., 2011). Breakpoints of conserved gene synteny between mammalian and chicken chromosomes were also mapped to the centromeres (International Chicken Genome Sequencing, 2004). Similar consequences in the karyotype have been reported in cases where centromeres were experimentally excised. Besides neocentromere formation, survival by fusion of acentric chromosome arms has been shown in *S. pombe* (Ishii et al., 2008). Such fusions are detected upon deletion of centromeres in another basidiomycete, *C. deuterogattii* (Schotanus and Heitman, 2019). By comparing the ancestral state (*M. slooffiae*) and other *Malassezia* species with either the same number or fewer chromosomes, we observed gene synteny breaks adjacent to centromeres (indicated by partial synteny conservation) apart from the break observed at *MgCEN2* or *MslCEN5*. Is this suggestive of a fragile nature of *Malassezia* centromeres? We advance the following hypothesis to explain the observed breaks at centromeres.

Studies of the common fragile sites in the human genome suggest different forms of replication stress as a major source of instability and subsequent breakage at these sites (Helmrich et al., 2011, Letessier et al., 2011, Ozeri-Galai et al., 2011). The resolution of the resulting replication fork stall has been shown to be critical for the stability of these fragile sites (Schwartz et al., 2005). Studies of the human fragile site FRA16D show that the AT-rich DNA (Flex1) results in fork stalling as a consequence of cruciform or secondary structure formation (Zhang and Freudenreich, 2007). Centromeres are natural replication fork stall sites in the genome (Greenfeder and Newlon, 1992, Smith et al., 1995, Mitra et al., 2014). The AT-rich core centromere sequence in *M. globosa* is also predicted to form secondary structures (Figure 7-figure supplement 4), which can be facilitated by the inherent replication fork stall at the centromeres. Whenever these secondary structures are unresolved and the fork restart fails, DSBs can occur at the centromeres. Chromosomal breakage and aneuploidy resulting from such defects are known in cancers (Kops et al., 2005). In mammals, centromeric DSBs are repaired efficiently compared to regions elsewhere in the genome, largely due to the presence of several homology tracts in the form of repetitive DNA sequences and the stiffness provided by the inherent heterochromatic state to facilitate ligation (Rief and Lobrich, 2002). *Malassezia* species are haploid in nature and lack typical pericentric heterochromatin marks. While the efficiency of centromeric DSB repair in the absence of long tracts of homologous sequences is not known in this species complex, we propose that the AT-rich core sequences, by virtue of secondary structure formation during DNA replication, could occasionally undergo DNA breakage at the centromere in *Malassezia* species.

The second mechanism of chromosome number reduction based on our comparative analyses of the *M. furfur* genome with genomes of *M. slooffiae* or *M. globosa* involves the inactivation of centromeres in the process of transition from a 9-chromosome state to a 7-chromosome state. Centromere inactivation occurs in cases involving the fusion of centric chromosomal fragments, stabilizing the fusion product and generating a functionally monocentric chromosome, as seen in case of the origin of human Chr2 from the shared ancestor with the great apes (Yunis and Prakash, 1982, Jdo et al., 1991). A larger proportion of known centromere inactivation events were shown to be mediated by epigenetic modification wherein inactivated centromeres are enriched with marks such as H3K9me2/3, H3K27me2/3, or DNA methylation, emphasizing the role of heterochromatin in this process (Zhang et al., 2010, Koo et al., 2011, Sato et al., 2012). Deletion of the centromere sequence corresponding to kinetochore binding has also been reported as an alternate mechanism, albeit less frequent, in both humans and in yeasts (Stimpson et al., 2010, Gordon et al., 2011, Sato et al., 2012). Unlike the above two modes, we observed divergence in the sequences corresponding to the inactivated centromeres (*CEN8* and *CEN9* of both *M. slooffiae* and *M. globosa*) in the arms of *M. furfur* Chr3, resulting in the loss of AT-richness of these centromere core regions (Figure 7C). This is also suggestive of a functional role for AT-rich DNA in centromere function in these species.

A change in chromosome number between two closely related species like *C. albicans* and *C. tropicalis* is associated with a change in centromere structure, between unique short regional centromeres in *C. albicans* that are epigenetically regulated, as compared to a genetically defined homogenized inverted repeat-associated centromere in *C. tropicalis* (Chatterjee et al., 2016). Strikingly, in both the transitions described above for *Malassezia* species, we did not observe any change in the centromere structure. The emergence of evolutionarily new centromeres as seen in the case of primates was not detected in *Malassezia* species (Rocchi et al., 2009, Kalitsis and Choo, 2012). This is particularly striking in the absence of conservation of any specific centromere-exclusive DNA sequence. This suggests that a strong driving force helps maintain the highly conserved centromere properties in closely related *Malassezia* species descended from their common ancestor, even after extensive chromosomal rearrangements involving centromeres that might have driven speciation. Furthermore, centromere inactivation/loss of centromere function seems to be a conserved theme mediating chromosome number variation from unicellular yeast species to metazoans, including primates.

## Materials and methods

The reagents, strains, plasmids, and primers used in this study are listed in the *SI appendix* (Tables S4-S7). *Malassezia* strains were grown on modified Dixon’s media (Malt extract 36 g/L, Desiccated Ox-bile 20 g/L, Tween40 10 mL/L, Peptone 6 g/L, Glycerol 2 mL/L, Oleic acid 2.89 mL/L). *M. sympodialis, M. furfur* strains were grown at 30°C. Cultures of *M. globosa* and *M. slooffiae* were grown at 32°C.

### Construction of the *M. sympodialis* strain expressing GFP-Mtw1

The allele for N-terminal tagging of Mtw1 with GFP was prepared by gap repair in the *Saccharomyces cerevisiae* BY4741 strain (Eckert-Boulet et al., 2012). Briefly, a 1.6 kb fragment consisting of the upstream and promoter sequence of the *MTW1* gene and a 1.6 kb fragment having the *MTW1* ORF (Protein ID: SHO76526) along with the downstream sequence was amplified from *M. sympodialis* genomic DNA. The GFP ORF (without the stop codon) and NAT were amplified from plasmids pVY7 and pAIM1 respectively (Yadav et al., 2018b). *S. cerevisiae* was transformed with all four fragments and the linearized plasmid pGI3 (digested with KpnI and BamHI) and the epitope-tagged allele were assembled in an ordered way by gap repair. Total DNA was isolated from *S. cerevisiae* and *E. coli* DH5α strain was transformed. The pGFP-Mtw1 construct was screened by restriction digestion and further confirmed by sequencing. The pGFP-Mtw1 construct was used to transform *M. sympodialis* strain ATCC42132 by *Agrobacterium tumefaciens*-mediated transconjugation (Ianiri et al., 2016, Ianiri et al., 2017a).

### Construction of the *M. furfur* strain expressing CENP-A FLAG

The allele for C-terminal tagging of CENP-A with a 3xFLAG epitope tag was prepared by gap repair in the *Saccharomyces cerevisiae* BY4741 strain (Eckert-Boulet et al., 2012). Briefly, a 1 kb fragment consisting of the upstream and promoter sequence of the CENP-A gene of *M. furfur* including the ORF (CENP-A ORF coordinates in Chr1: 1,453,468-1,453,921) and 1 kb fragment containing the sequence downstream of CENP-A ORF were amplified from *M. furfur* genomic DNA. The 3xFLAG tag was introduced in the reverse primer annealing to the CENP-A ORF. The NAT marker was amplified from plasmid pAIM1 as above. *S. cerevisiae* was transformed with all three fragments and plasmid pGI3 (digested with KpnI and BamHI) and the epitope-tagged allele was assembled in an ordered way by gap repair. Total DNA was isolated from *S. cerevisiae* and *E. coli* DH5α strain was transformed. The resulting pMF1 construct was screened by restriction digestion and further confirmed by sequencing. The pMF1 construct was used to transform *M. furfur* strain CBS14141 by *Agrobacterium tumefaciens*-mediated transconjugation (Ianiri et al., 2016, Ianiri et al., 2017a) to obtain the epitope-tagged strain MF001.

### Microscopic imaging of live cells and processing

The GFP-Mtw1 strain was inoculated to 1% v/v from a saturated starter culture grown in mDixon medium. After growth for 6 h at 30°C/ 150 rpm, these cells were pelleted at 4,000 rpm and washed 3 times with 1x phosphate-buffered saline (PBS) and the cell suspension was placed on a clean glass slide. A coverslip was placed on the spot and sealed prior to imaging. The images were acquired at room temperature using a laser scanning inverted confocal microscope LSM 880-Airyscan (ZEISS, Plan Apochromat 63x, NA oil 1.4) equipped with highly sensitive photo-detectors. The filters used were GFP/FITC 488 excitation and GFP/FITC 500/550 band pass, long pass for emission. Z-stack images were taken at every 0.3 μm and processed using ZEISS Zen software or ImageJ. All of the images were processed post-acquisition with minimal adjustments to levels and linear contrast until the signals were highlighted.

### Preparation of spheroplasts

Cells were grown on mDixon’s medium and washed with water by centrifugation at 4000 rpm for 5 min. Cells were resuspended in 10 mL of 5% (v/v) 2-mercaptoethanol solution in water and incubated at 30°C/ 150 rpm for 45 min. The cells were pelleted, washed, and resuspended in 3 mL spheroplasting buffer (40 mM Citric acid, 120 mM Na_2_HPO_4_ and 1.2 M Sorbitol) for every 1.5×10^9^ cells. Cell clumps were dissociated by mild sonication for 30 s using the medium intensity setting in a Bioruptor (Diagenode). Lysing enzymes from *Trichoderma harzianum* (Sigma) and Zymolyase-20T (MP Biomedicals) were added at 20 mg/mL and 100 µg/mL respectively. The spheroplasting suspension was incubated at 30°C/ 65 rpm for 6 to 8 h. The suspension was examined under a microscope to estimate the proportion of spheroplasts. Spheroplasts were washed with ice-cold 1xPBS and used as per the experimental design (adapted from (Boekhout, 2003)).

### Indirect Immunofluorescence

The GFP-Mtw1 strain was inoculated to 1% (v/v) from a saturated starter culture grown in mDixon medium. After growth for 6 h, the cells were fixed by the addition of formaldehyde to a final concentration of 3.7% for 1 h. Post-fixing, the cells were washed with water and taken for preparation of spheroplasts (described above). Spheroplasts were washed with ice-cold 1xPBS and resuspended in ice-cold 1xPBS to a cell density suitable for microscopy. Slides for microscopy were washed with water and coated with poly L-Lysine (15 µL of 10 mg/mL solution per well) for 5 min at room temperature. The solution was aspirated and washed once with water. The cell suspension was added to each well (15-20 µL) and allowed to stand at room temperature for 5 min. The cell suspension was aspirated and the slides were washed once with water to remove unbound cells. The slides were fixed in ice-cold methanol for 6 min followed by treatment with ice-cold acetone for 30 s. Post fixing, blocking solution (2% non-fat skim milk in 1xPBS) was added to each well, incubated at room temperature for 30 min. After this, the blocking solution was aspirated and primary antibodies were added (mouse anti-GFP antibodies [Sigma] at 1:100 dilution). After incubation for 1 h at room temperature, each slide was washed 8 times with 1xPBS giving a 2 min incubation for every wash. Secondary antibody solution (goat anti-mouse-AlexaFluor488 [Invitrogen] at 1:500 dilution) was added to each well and incubated for 1 h in the dark at room temperature. Post-incubation, slides were washed as described above. Mounting medium (DAPI at 100 ng/mL in 70% glycerol) was added, incubated for 5 min and aspirated out. Slides were sealed with a clean coverslip and proceeded for imaging. The images were acquired at room temperature using an inverted fluorescence microscope (ZEISS Axio Observer, Plan Apochromat 100x, NA oil 1.4). Z-stack images were taken at every 0.3 μm and processed using ZEISS Zen software/ ImageJ.

### Chromatin immunoprecipitation (ChIP)

The ChIP protocol was adapted from the one implemented for *C. neoformans* (Yadav et al., 2018b). Logarithmically grown cells were fixed with formaldehyde at a final concentration of 1% for 30 min (for Mtw1 ChIP) and 15 min (for CENP-A, histone H3, and histone H4 ChIP) respectively. The reaction was quenched by the addition of glycine to a final concentration of 0.135 M. Cells were pelleted and processed for spheroplasting as described above. Spheroplasts were washed once sequentially using 10 mL of the following ice-cold buffers: 1xPBS, Buffer-I (0.25% Triton X-100, 10 mM EDTA, 0.5 mM EGTA, 10 mM Na-HEPES pH=6.5), Buffer-II (200 mM NaCl, 1 mM EDTA, 0.5 mM EGTA, 10 mM Na-HEPES pH=6.5). The pellet after the final wash was resuspended in 1 mL lysis buffer (50 mM HEPES pH=7.4, 1% Triton X-100, 140 mM NaCl, 0.1% Na-deoxycholate, 1 mM EDTA) for every 1.5×10^9^ cells. Protease inhibitor cocktail was added to 1x final concentration.

The resuspended spheroplasts were sonicated with a Bioruptor (Diagenode) using 30 s ON/OFF pulse at high-intensity mode with intermittent incubation on ice to obtain chromatin fragments in the size range of 100-300 bp. The lysate was cleared after sonication by centrifugation at 13,000 rpm for 10 min at 4°C. The input DNA fraction was separated at this step (1/10^th^ volume of lysate) and processed for de-crosslinking by the addition of 400 µL elution buffer (0.1 M NaHCO_3_, 1% SDS) per 100 µL lysate (processing for de-crosslinking mentioned below). The remaining lysate was split equally and processed as IP and control samples. For GFP-Mtw1 ChIP, 20 µL GFP-trap beads and blocked agarose beads respectively were used for IP and control. For CENP-A-3xFLAG ChIP, 20 µL anti-FLAG affinity gel and blocked agarose beads respectively were used for IP and control. In the case of histone H3 or histone H4 ChIP, 5 µL of antibodies were used per IP fraction along with 20 µL Protein-A sepharose beads. Samples were rotated for 6 h at 4°C. Post incubation, samples were sequentially washed as follows: twice with 1 mL low salt wash buffer (0.1% SDS, 1% Triton X-100, 2 mM EDTA, 20 mM Tris pH=8.0, 150 mM NaCl), twice with 1 mL high salt wash buffer (0.1% SDS, 1% Triton X-100, 2 mM EDTA, 20 mM Tris pH=8.0, 500 mM NaCl), once with 1 mL LiCl wash buffer (0.25 M LiCl, 1% NP-40, 1% Na-deoxycholate, 1 mM EDTA, 10 mM Tris pH=8.0) and twice with 1 mL 1xTE (10 mM Tris pH=8.0, 1 mM EDTA). Samples were rotated in a rotaspin for 5 min at room temperature for every wash (15 min in case of histone H3 or histone H4 ChIP). After washes, DNA was eluted from the beads twice using 250 µL elution buffer. The samples for elution were incubated at 65°C for 5 min, rotated for 15 min and then collected by centrifugation.

Samples were decrosslinked by the addition of 20 µL 5 M NaCl and incubation at 65°C for 6 h. Following this, samples were deproteinized by the addition of 10 µL 0.5 M EDTA, 20 µL 1 M Tris pH=6.8, 2 µL Proteinase K (20 mg/L) and incubation at 45°C for 2 h. After incubation, samples were treated with an equal volume of phenol-chloroform-isoamyl alcohol (25:24:1) mix, and the aqueous phase was extracted by centrifugation. DNA was precipitated by addition of 1/10^th^ volume of 3 M Na-acetate, 1 µL glycogen (20 mg/mL), 1 mL absolute ethanol and incubation at −20°C for at least 12 h. Finally, the samples were harvested by centrifugation at 13,000 rpm for 45 min at 4°C followed by washing the pellet once with ice-cold 70% ethanol. Air-dried pellets were then resuspended in 20 µL sterile MilliQ water with 10 µg/mL RNAse. Samples were either processed for library preparation for ChIP-sequencing or analyzed by qPCR with the primers listed in Table S7.

### Analysis of sequencing data

GFP-Mtw1 ChIP sequencing was performed at the Clevergene Biocorp. Pvt. Ltd., Bengaluru, India. A total of 46,704,720 and 63,524,912 150 bp paired-end reads were obtained for IP and Input samples, respectively. The reads were mapped to *M. sympodialis* ATCC42132 genome using Geneious 9.0 (http://www.geneious.com/) with default conditions. Each read was allowed to map only once randomly anywhere in the genome. The alignments were exported to BAM files, sorted, and visualized using Integrative Genomics Viewer (IGV, Broad Institute). The images from IGV were imported into Adobe Photoshop (Adobe) and scaled for representation purposes. RNA-sequencing data (E-MTAB-4589) from a previous study was downloaded from the ArrayExpress website, sorted, and visualized using IGV. GC-content was calculated using Geneious 9.0 with a sliding window size of 250 bp. The data was exported as wig files and further visualized using IGV.

### Western Blotting

Protein lysates for western blot were prepared by the TCA method. One mL overnight grown cultures were harvested, washed, and resuspended in 400 μL of 12.5% ice-cold TCA solution. The suspension was vortexed briefly and stored at – 20°C for 4 to 6 h. The suspension was thawed on ice, pelleted at 14,000 rpm for 10 min, and washed twice with 350 μL of 80% Acetone (ice-cold). The washed pellets were air-dried completely and resuspended in the desired volume of lysis buffer (0.1 N NaOH+1% SDS). Samples were separated in 12% polyacrylamide gels and transferred onto nitrocellulose membranes. For probing, mouse anti-GFP antibody (Roche) and the HRP conjugated goat anti-mouse secondary antibody (Bangalore Genei), were used at 1:3000 and 1:5000 dilution respectively in 2.5% skim milk powder in 1xPBS. The blots were developed using Chemiluminescence Ultra substrate (BioRad) and imaged using the VersaDoc imaging system (BioRad).

### PFGE analysis for *M. globosa* and *M. slooffiae*

For CHEF analysis of *M. globosa* (CBS7966) and *M. slooffiae* (CBS7956), the cells were grown on solid mDixon medium and then collected and resuspended in 1xPBS. CHEF plugs were prepared as described in previous studies (Sun et al., 2014, Sun et al., 2017). Chromosomes were separated in 1% Megabase certified agarose gels made with 0.5×TBE, using a BioRad CHEF-DR II System running at 3.2 V/cm with linear ramping switching time from 90 to 360 s for 120 h in 0.5xTBE at 14°C. The gel was stained with EtBr and visualized under UV.

For the chromoblot analyses of *M. globosa* (CBS7966), the gel from the CHEF analysis was first transferred to a membrane, and the resulting chromoblots were then hybridized sequentially with four probes from the chromosomes 3, 4, 5, and 6 of the CBS7966 genome assembly, respectively (see the Table S7 for the primer information), as described in previous studies (Findley et al., 2012, Yadav et al., 2018b).

### *M. globosa* genome assembly

Sequence reads were assembled with HGAP3 included in SMRTPortal v2.3 (PacBio, Menlo Park, CA, USA) and default parameters except for the genome size set to 9 Mb. Assembly completeness was evaluated by checking for telomeric repeats. Non-telomeric contig-ends were aligned to other contigs using BLAST and unique overlaps used to build complete chromosomes. Short telomere ends were extended using uniquely mapping reads longer than 10 kb and repolishing of the assembly using the resequencing pipeline in SMRTPortal v2.3. The assembly resulted in 19 contigs, with a total length of 9.2 Mb. 17 long and 1 short telomere could be identified. 6 contigs had telomeres on both the 5’- and the 3’-end and thus represent full-length chromosomes (Chr1, 2, 3, 6, 7, and 8). 6 contigs had only one telomere and 7 contigs had no telomeric sequence. 2 contigs without telomeres were from the mitochondrion and 2 were from the ribosomal repeats. Chromosome 5 was constructed from 2 contigs ending in ribosomal rDNA repeats. The assembly contains 6 copies of the repeat, but read coverage suggests a length of 30-40 repeat units that cannot be resolved with the available read-length. The remaining contigs were used to build chromosomes 4 and 9 that share highly similar 5’-ends. The two ends can be distinguished by two microsatellite expansions. Chromosome 4 had a very short 3’-telomere from the default assembly, but the raw data contained a uniquely mapping read that extended several repeat units past the assembly end. After polishing the reference, all 9 chromosomes had clear 5’- and 3’-telomeres.

### *M. slooffiae* genome assembly

Sequence reads were assembled using HGAP3 included in SMRTPortal v2.3 (PacBio, Menlo Park, CA, USA) with default parameters. This resulted in an assembly with 14 scaffolds with telomeric repeats at both ends in 9 contigs. Of the remaining five contigs, three of them could be assigned to mitochondrial DNA based on BLAST analysis with *M. globosa.* The remaining two contigs of sized 5.8 kb and 2.3 kb respectively did not show BLAST hits against *M. globosa* or *M. sympodialis* genomes.

### *M. furfur* genome assembly

Sequence reads generated from CBS14141 were assembled using HGAP3 included in SMRTPortal v2.3 (PacBio, Menlo Park, CA, USA) with default parameters. This resulted in an assembly with 8 scaffolds of which the nuclear genome was organized in 7 scaffolds having telomere repeats at both ends.

### Gene synteny analysis

For gene synteny conservation across the centromeres, the analysis was performed by BLAST as follows. The genomes for *M. restricta*, *M. nana,* and *M. dermatis* were downloaded from the NCBI genomes portal. The PacBio assembled genomes of *M. globosa* and *M. slooffiae* were used for synteny analysis. Synteny analysis was done in the context of ORFs flanking the centromeres of *M. sympodialis*. The protein sequences for each of these ORFs served as the query in BLAST analysis against the genome of other species. The local database for each genome was set up in the Geneious software for this analysis. The percentage identity values for each ORFs are mentioned in the boxes in Figure 7-figure supplement 3. Additionally, synteny analyses between *M. globosa* and *M. sympodialis* were conducted with megablast (word size: 28) and plotted together with GC content (calculated as the deviation from the genomic mean, in non-overlapping 1 kb windows), using Circos (v0.69-6) (Krzywinski et al., 2009). Additional whole-genome alignments were conducted with Satsuma (Grabherr et al., 2010), with default parameters. The linear synteny comparisons shown in Figures 5 and 6 were generated with the Python application EasyFig (Sullivan et al., 2011).

### Species phylogeny

To reconstruct the phylogenetic relationship among the 9 *Malassezia* species selected, orthologs were identified using the bidirectional best-hit (BDBH), COGtriangles (v2.1), and OrthoMCL (v1.4) algorithms implemented in the GET_HOMOLOGUES software package (Contreras-Moreira and Vinuesa, 2013). The proteome of *M. sympodialis* ATCC42132 was used as a reference. A phylogeny was inferred from a set of 738 protein sequences as follows. Individual proteins were aligned using MAFFT v7.310 (L-INS-i strategy) and poorly aligned regions were trimmed with TrimAl (-gappyout). The resulting alignments were concatenated to obtain a final supermatrix consisting of a total of 441,200 amino acid sites (159,270 parsimony-informative). This sequence was input to IQ-TREE v1.6.5 (Nguyen et al., 2015) and a maximum likelihood phylogeny was estimated using the LG+F+R4 amino acid model of substitution. Branch support values were obtained from 10,000 replicates of both ultrafast bootstrap approximation (UFBoot) and the nonparametric variant of the approximate likelihood ratio test (SH-aLRT) implemented in IQ-TREE. The best likelihood tree was graphically visualized with iTOL v4.3.3 (Letunic and Bork, 2019).

## Data availability

The Mtw1 ChIP sequencing reads reported in this paper have been deposited under NCBI BioProject (Accession number PRJNA509412). The genome sequence assemblies of *M. globosa*, *M. slooffiae,* and *M. furfur* have been deposited in GenBank with accession numbers SAMN10720087, SAMN10720088, and SAMN13341476 respectively.

## Acknowledgments

We thank Clevergene Biocorp Pvt. Ltd., Bengaluru for generating the Mtw1 ChIP-sequencing data. S.R.S is a Senior Research Fellow supported by intramural funding from JNCASR. M.H.R is a National Postdoctoral Fellow (PDF/2016/002858), supported by the Science and Engineering Research Board (SERB), Department of Science and Technology (DST), Government of India. K.S is a Tata Innovation Fellow (grant number BT/HRT/35/01/03/2017) and is supported by a grant for Life Science Research, Education and Training (BT/INF/22/SP27679/2018) of Department of Biotechnology, Govt. of India and intramural funding from JNCASR. Studies were supported in part by NIH/NIAID R37 award AI39115-21 and R01 award AI50113-15 to J.H. J.H is co-director and fellow of CIFAR program Fungal Kingdom: Threats & opportunities. T.L.D acknowledges the A* STAR Industry alignment fund H18/01a0/016, Asian Skin Microbiome Program. We thank the members of K.S lab and J.H lab for valuable discussions and comments during bi-weekly Skype meetings.

## Author contributions

K.S and J.H conceived the study and secured funding. S.S performed PFGE and chromoblot analysis of *M. slooffiae* and *M. globosa*. M.H.R generated the constructs for tagging MsyMtw1 and MfCENP-A. Transformation of *Malassezia* strains and PCR confirmation was done by G.I. S.R.S performed western blot, microscopy, ChIP-qPCR analysis, and prepared the samples for ChIP-sequencing. P.G analyzed the ChIP-seq data and RNA-seq data for *M. sympodialis*. S.R.S, B.C.T, M.H.R, and M.A.C performed the synteny conservation analysis. M.A.C performed genome comparisons, phylogenetic analysis, and prepared the figures. R.N.V and R.S generated the GC/GC3 plots and performed motif analysis reported in this study. T.L.D and C.T.R provided the genome assemblies of *M. globosa*, *M. slooffiae*, and *M. furfur*. S.R.S and K.S wrote the manuscript with help from all of the authors. All of the authors revised and approved the final version.

## Figures and figure supplements

**Figure 1-figure supplement 1.**
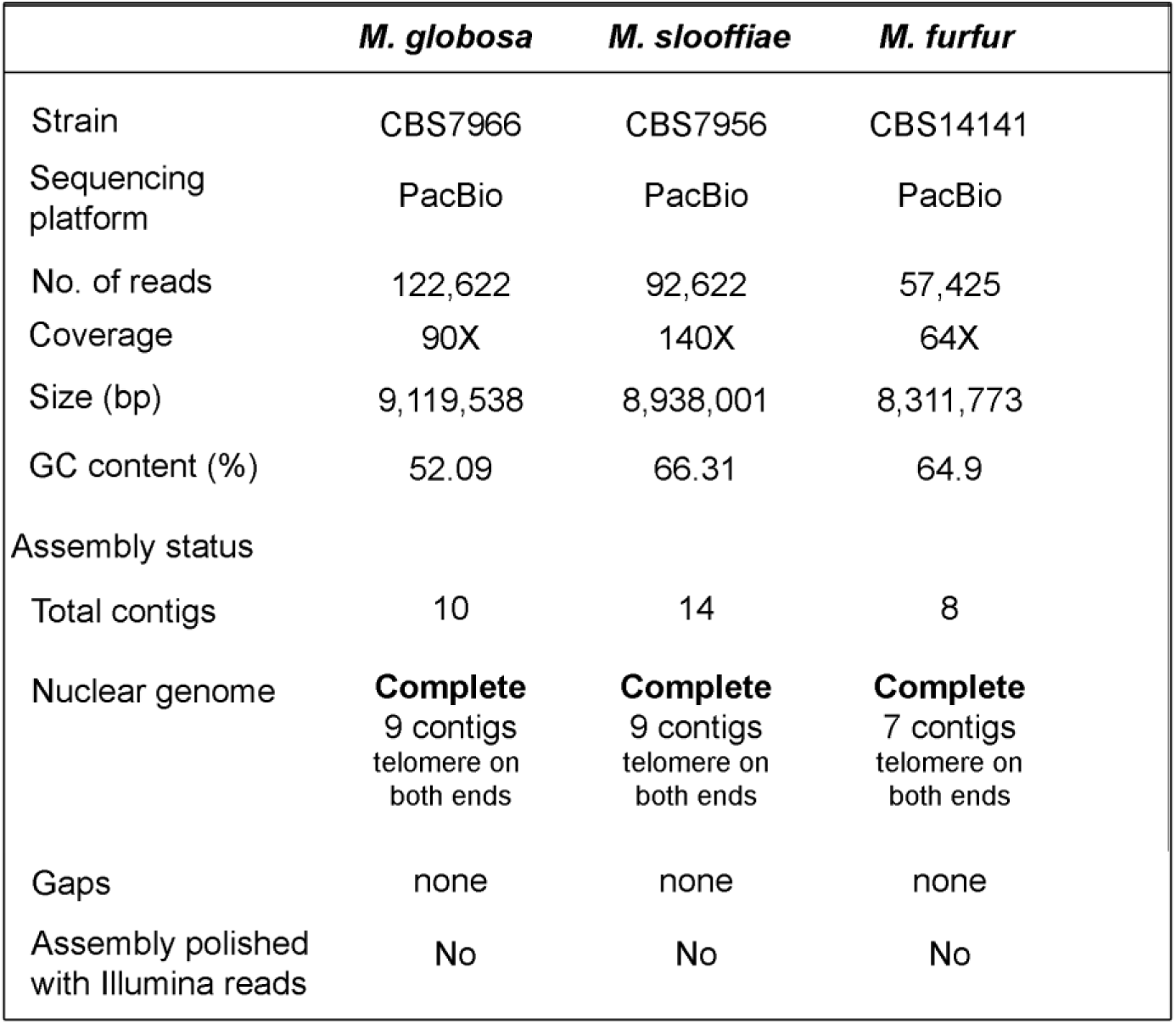
Statistics of the genome assemblies of *M. globosa*, *M. slooffiae,* and *M. furfur* generated in this study.

**Figure 2-figure supplement 1.**
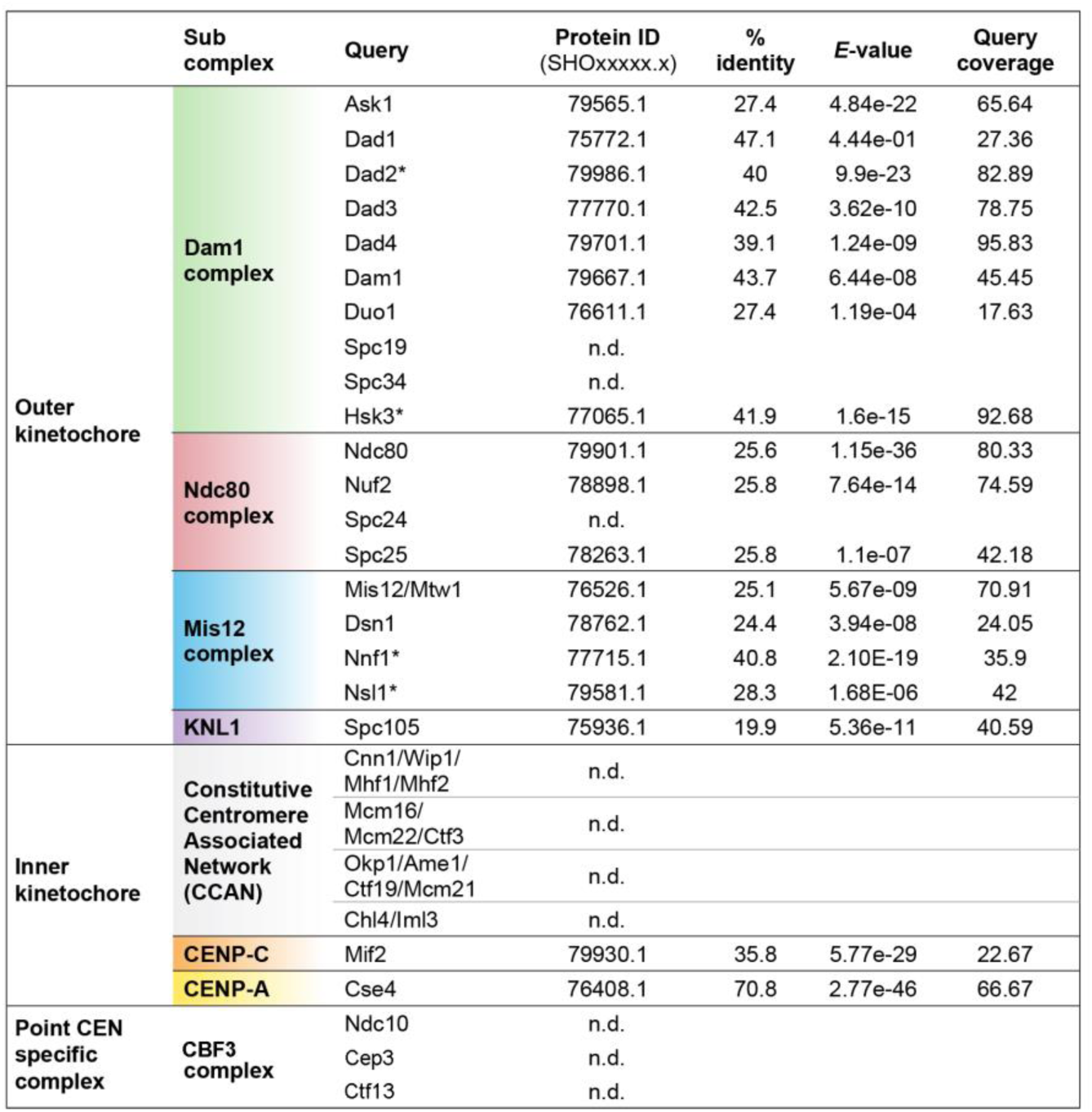
Identification of kinetochore proteins in *M. sympodialis* by BLAST. Putative homologs of the indicated proteins in *M. sympodialis* were identified by BLAST analysis using corresponding protein sequences of *C. neoformans* as the query. Asterisk (*) indicates cases where corresponding protein sequences from *Ustilago maydis* were used as query. To detect proteins of the CBF3 complex, protein sequences of Ndc10, Cep3, and Ctf13 from *S. cerevisiae* were used as query. ‘n.d.’ indicates homolog not detected.

**Figure 3-figure supplement 1.**
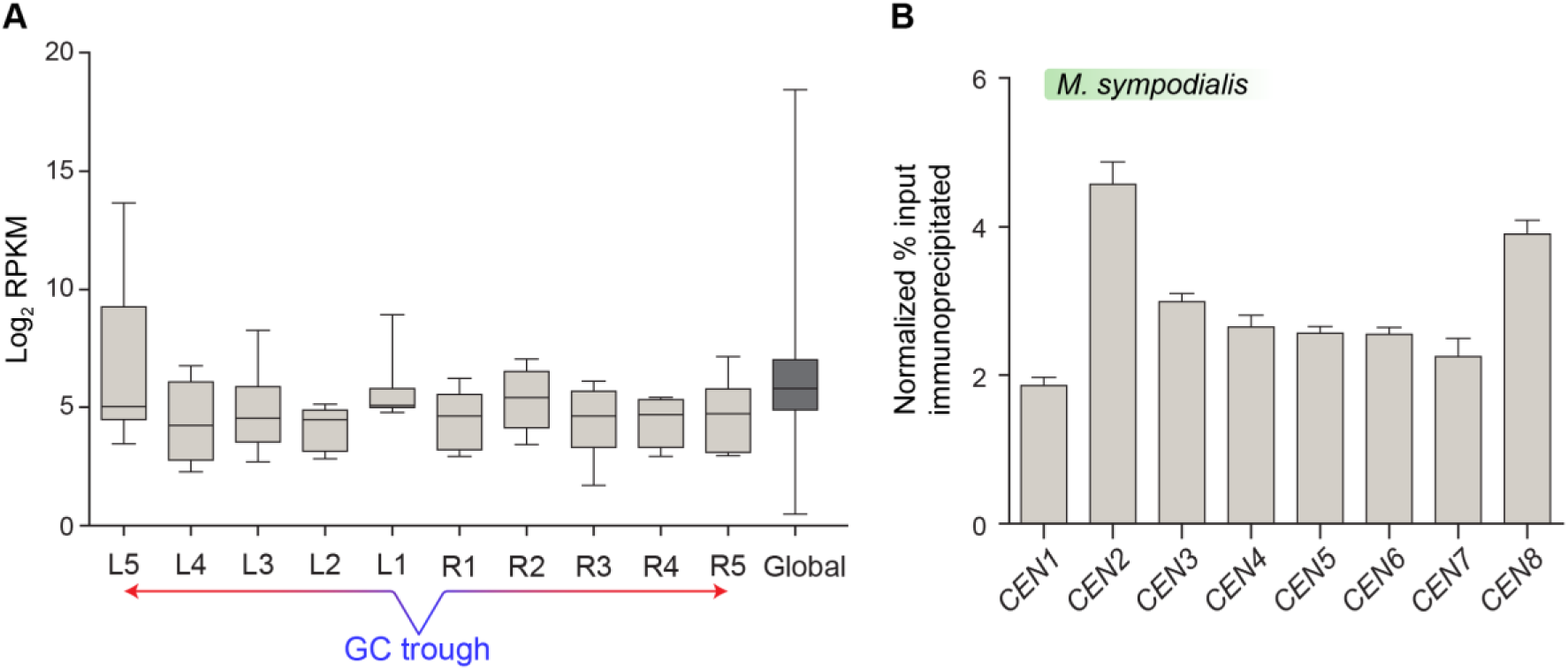
Mtw1-enriched regions in *M. sympodialis* contain transcriptionally active genes. (A) Normalized read counts (RPKM) for transcripts from ORFs flanking the centromere as compared with the genome average RPKM value calculated from RNA-seq data of *M. sympodialis* (ATCC42132). The *x*-axis represents ORFs labeled L1 - L5 and R1 - R5, with L1 and R1 being proximal to the GC trough of the centromere. Genome average considered for comparison is labeled as global. The *y*-axis represents Log_2_ RPKM values. (B) ChIP-qPCR assays validating the enrichment of Mtw1 at the centromeres identified by ChIP-sequencing analysis. The *x*-axis indicates individual *CEN* regions assayed with primers mapping to the chromosomal GC troughs (see Table S7 for primer sequences) and the *y*-axis indicates enrichment of Mtw1 over a centromere unlinked control region as ‘normalized % input immunoprecipitated’. Error bars indicate standard deviation (SD).

**Figure 3-figure supplement 2.**
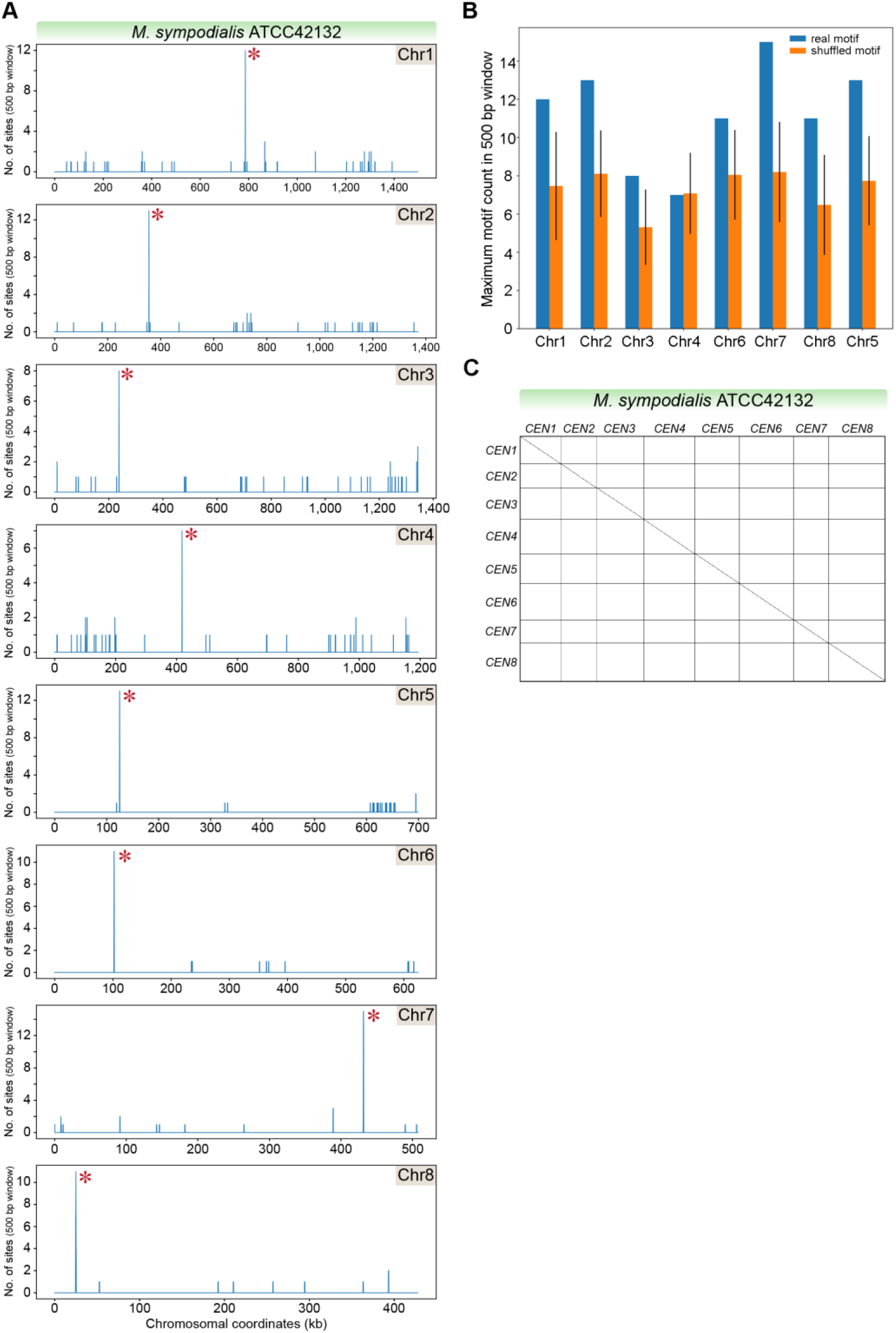
Sequence features of centromeres in *M. sympodialis.* (A) The genome of *M. sympodialis* was scanned for matches to the 12 bp AT-rich motif using a 500 bp sliding window. Hit counts (*y*-axis) were plotted against the chromosomal coordinates (*x*-axis, in kb). Red asterisks near the line corresponding to maximum enrichment in every chromosome mark the regions predicted as centromeres. (B) Comparison of the maximum motif counts in a 500 bp window in each *M. sympodialis* chromosome (*y-*axis) estimated with the PWM of the 12 bp motif (blue bars, labelled real) and a scrambled version of the matrix (orange bars, labelled scrambled). The *x-*axis indicates individual chromosomes in *M. sympodialis* (C) Dot-plot generated by plotting sequences that include 5 kb flanking the Mtw1-enriched regions in *M. sympodialis* against themselves is depicted. The coordinates of regions used in *M. sympodialis* are *CEN1*: 781716-791716, *CEN2*: 350852-360852, *CEN3*: 232777-242777, *CEN4*: 413320-423320, *CEN5*: 120251-130251, *CEN6*: 97796-107796, *CEN7*: 426611-436611 and *CEN8*: 19905-29905.

**Figure 3-figure supplement 3.**
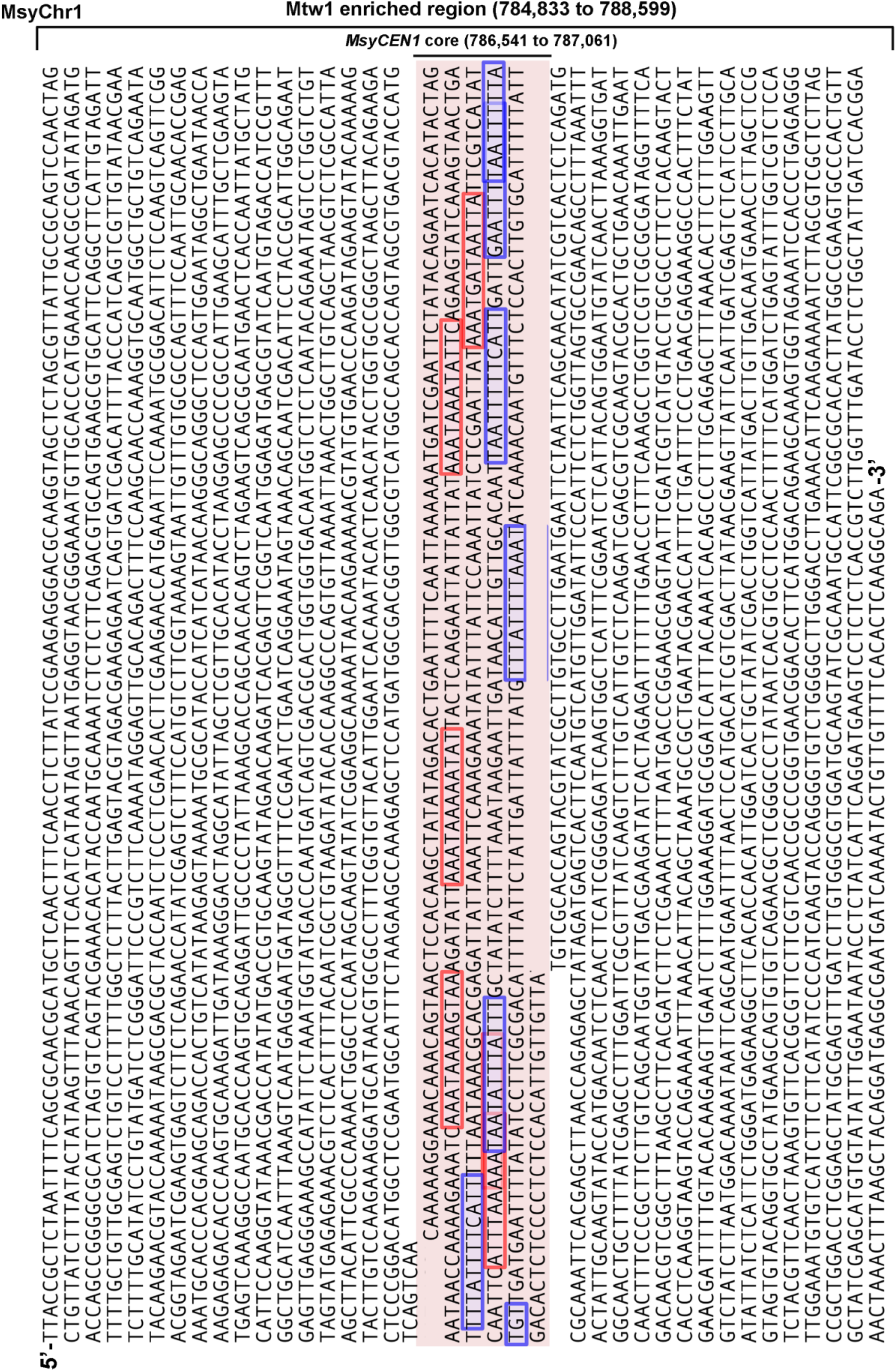
Enrichment of the 12 bp motif at the centromere core in *M. sympodialis*. The sequences corresponding to the Mtw1 enriched regions in Chr1 (with the AT-rich core region highlighted in pink) are taken as a representative of *M. sympodialis* centromeres. The sequences corresponding to the 12 bp motif shown in figure 3D are highlighted. Red and blue highlights indicate the forward and reverse orientation of motif occurrence.

**Figure 4-figure supplement 1.**
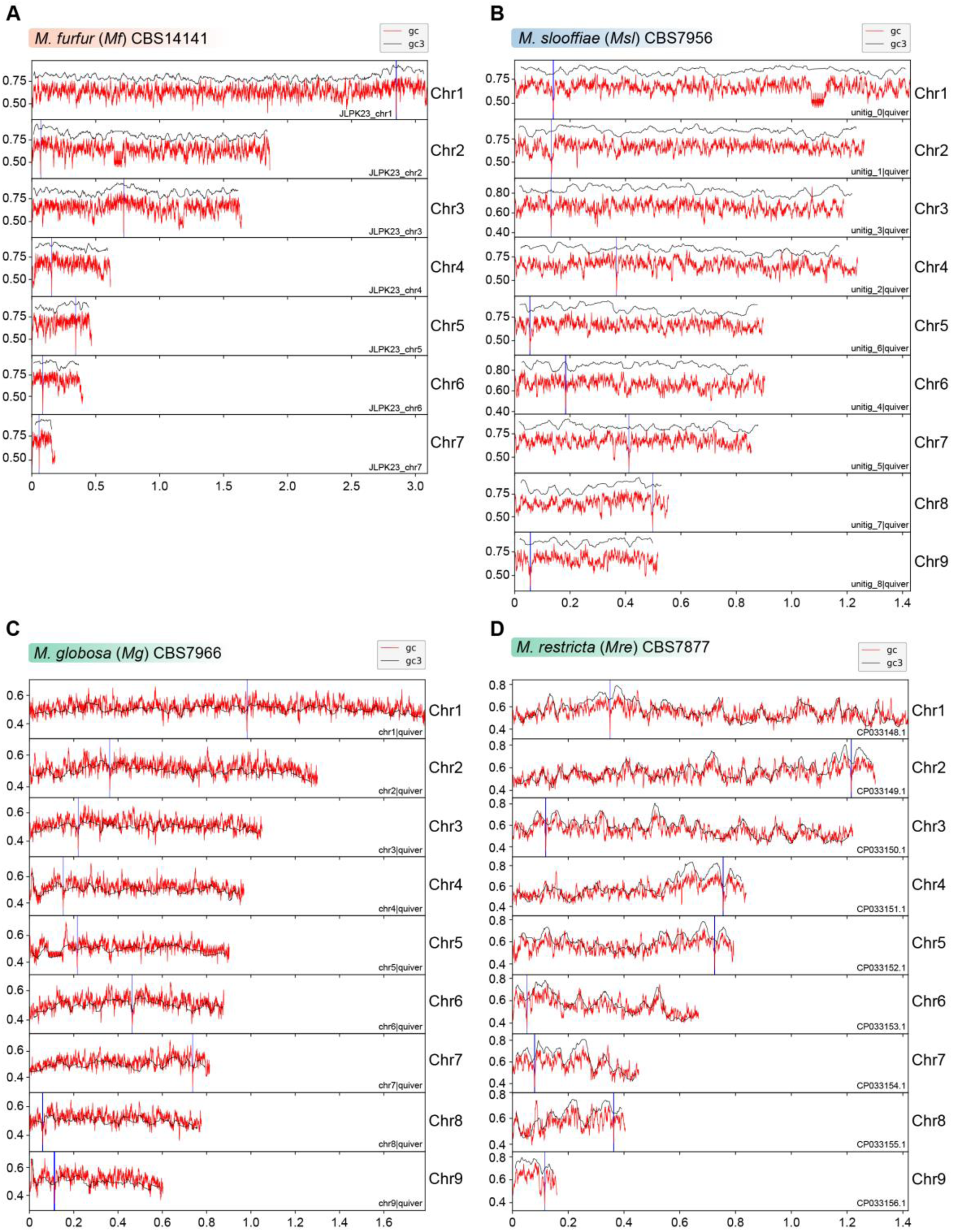
Putative centromeres of *M. furfur*, *M. slooffiae*, *M. globosa,* and *M. restricta* map to global GC troughs in each chromosome. (A-D) Graphs indicating GC content (red lines) and GC3 content (black lines) of each chromosome of *M. furfur*, *M. slooffiae, M. globosa,* and *M. restricta.* Position of putative centromeres mapping to chromosomal GC minima are marked in blue. The *x*-axis indicates chromosomal coordinates in Mb.

**Figure 4-figure supplement 2.**
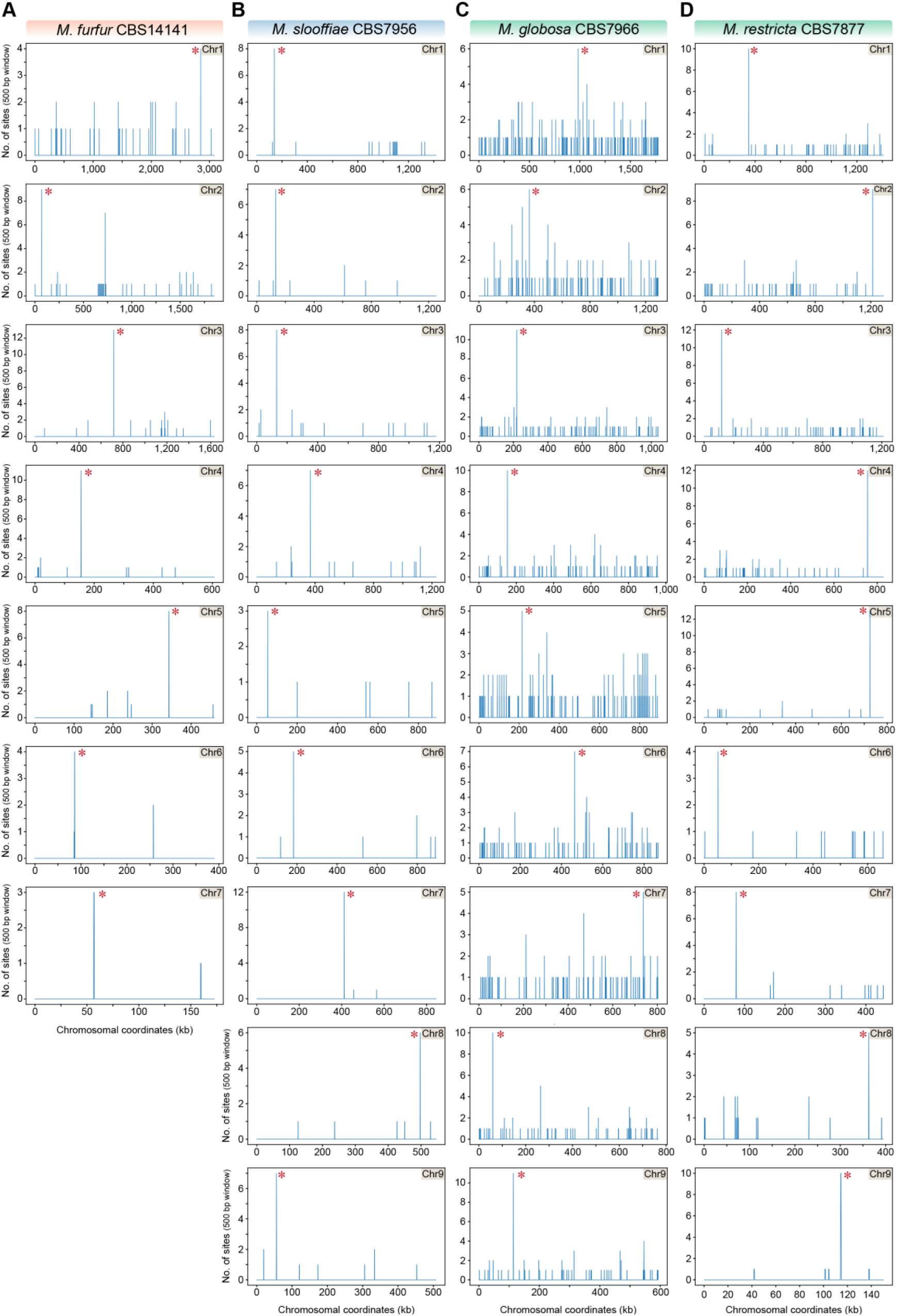
The 12 bp AT-rich motif is enriched at the putative centromeres of *M. furfur, M. slooffiae, M. globosa,* and *M. restricta*. (A-D) The genomes of *M. furfur, M. slooffiae, M. globosa,* and *M. restricta* were scanned for matches to the 12 bp AT-rich motif using a 500 bp sliding window. Hit counts (*y-*axis) were plotted against the chromosomal coordinates (*x-*axis, in kb) for each of the above species. Red asterisks near the line corresponding to maximum enrichment in every chromosome or scaffold mark the regions predicted as centromeres in each species.

**Figure 6-figure supplement 1.**
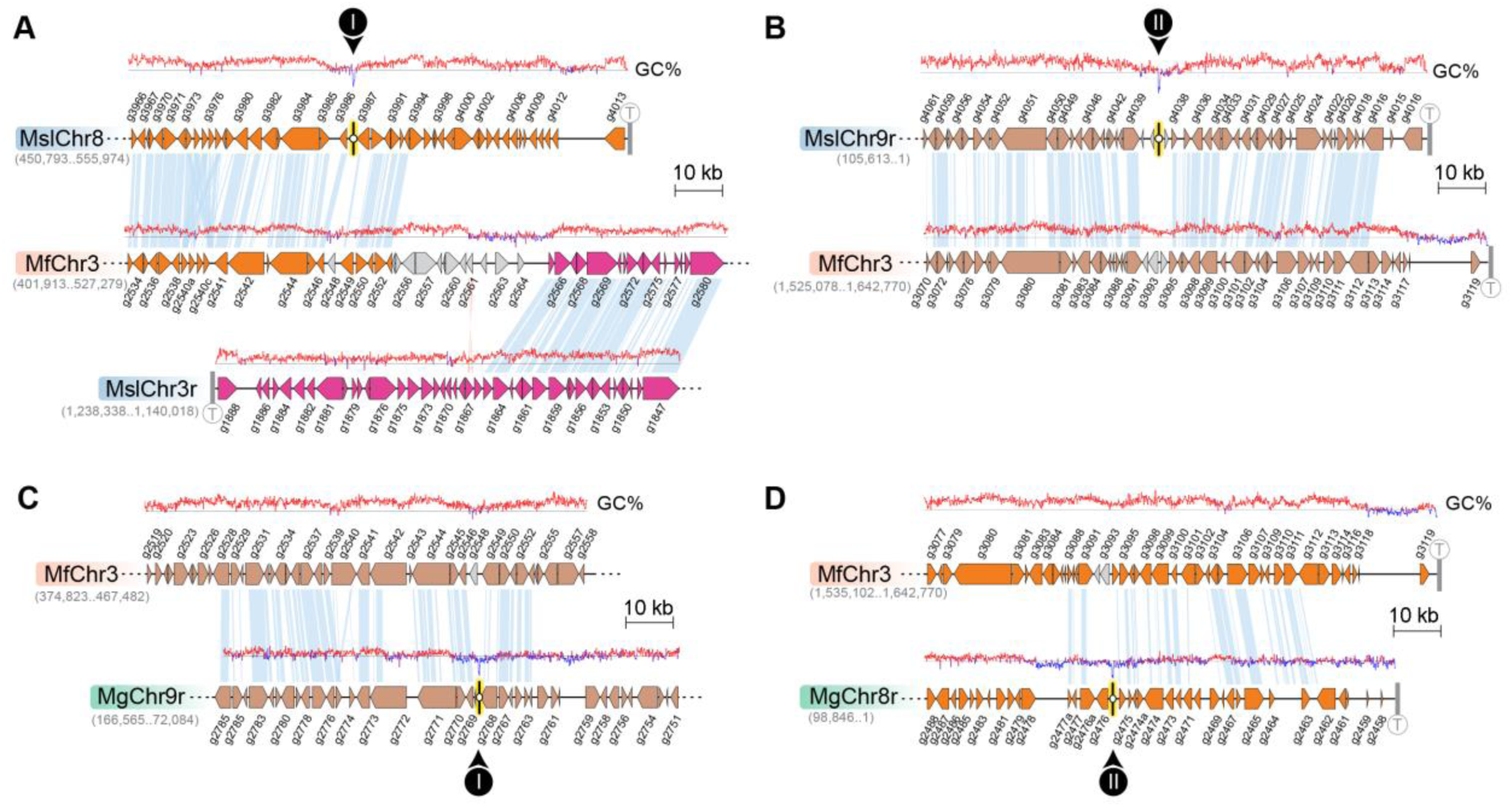
Loss of AT-content in regions corresponding to *CEN8* and *CEN9* of *M. slooffiae* and *M. globosa* in MfChr3. (A, B) Zoomed-in view of regions marked I and II in figure 6B, corresponding to inactivated *MslCEN8* and *MslCEN9* in *M. furfur* along with the regions of gene synteny is represented. GC content (in %) is shown as red/blue line diagram above each chromosome. (C, D) Zoomed-in view of the regions marked I and II in figure 6D, corresponding to inactivated *MgCEN9* and *MgCEN8* in *M. furfur* along with the regions of gene synteny is represented. GC content (in %) is shown as red/blue lines above each chromosome.

**Figure 7-figure supplement 1.**
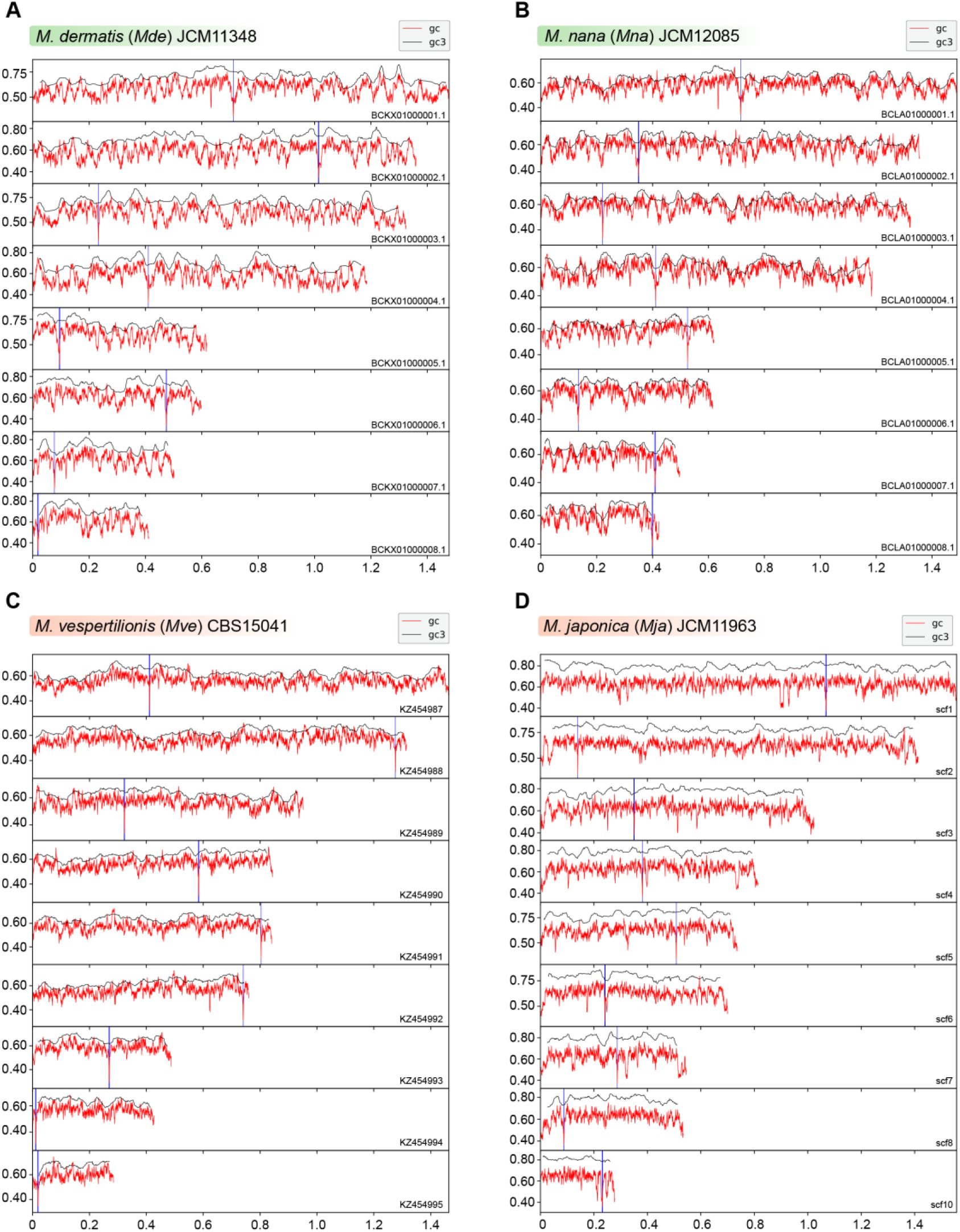
Putative centromeres of *M. dermatis*, *M. nana*, *M. vespertilionis,* and *M. japonica* map to global GC troughs in each chromosome. (A-D) Graphs indicating GC content (red lines) and GC3 content (black lines) of each chromosome of *M. dermatis*, *M. nana*, *M. vespertilionis,* and *M. japonica.* Position of putative centromeres mapping to chromosomal GC minima are marked in blue. The *x-*axis indicates chromosomal coordinates in Mb.

**Figure 7-figure supplement 2.**
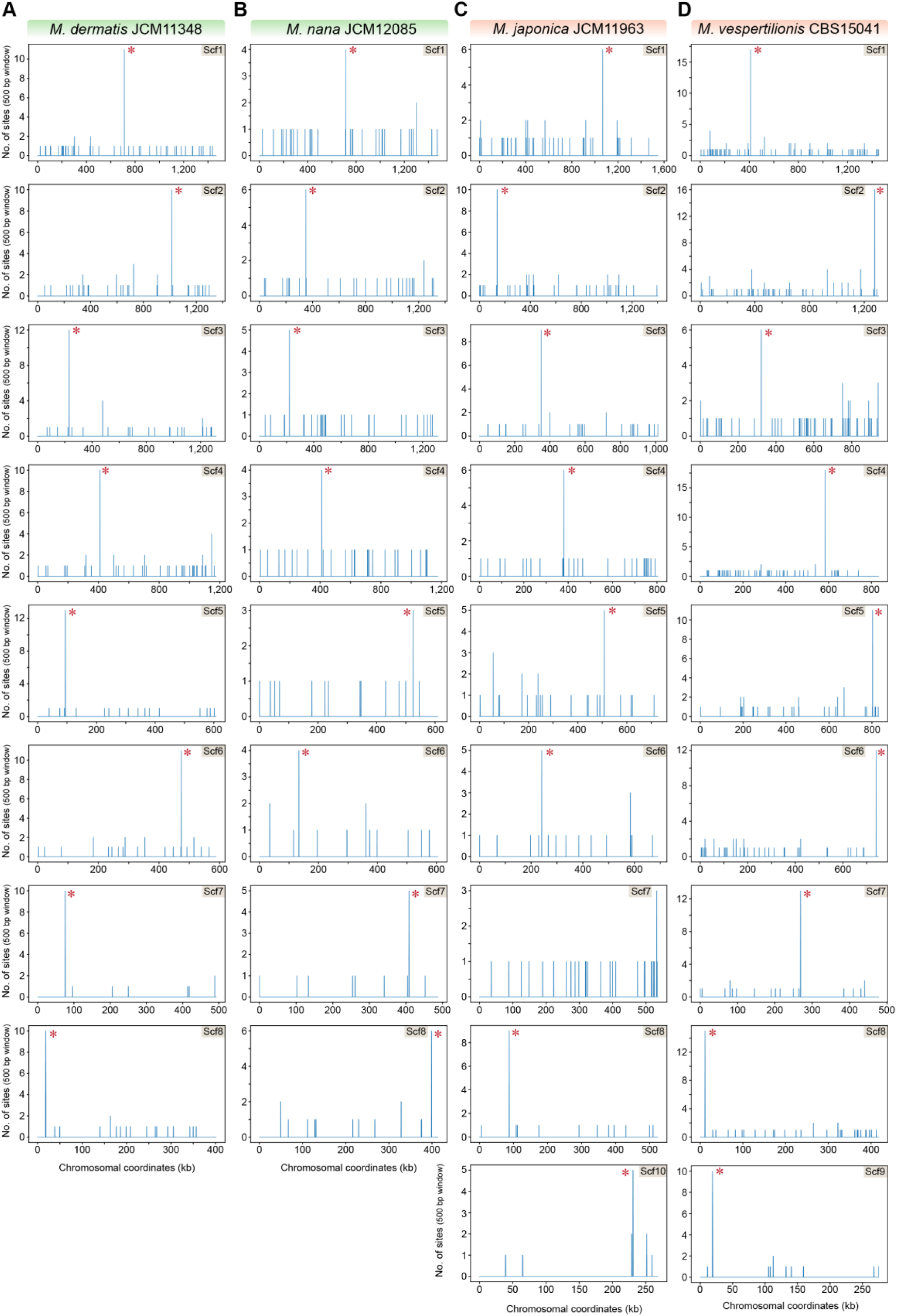
The 12 bp AT-rich motif is enriched at the putative centromeres of *M. dermatis, M. nana, M. japonica,* and *M. vespertilionis*. The genomes of *M. dermatis, M. nana, M. japonica,* and *M. vespertilionis* were scanned for matches to the 12 bp AT-rich motif using a 500 bp sliding window. Hit counts (*y-*axis) were plotted against the chromosomal coordinates (*x-*axis, in kb) for each of the above species. Red asterisks near the line corresponding to maximum enrichment in every chromosome or scaffold mark the regions predicted as centromeres in each species.

**Figure 7-figure supplement 3.**
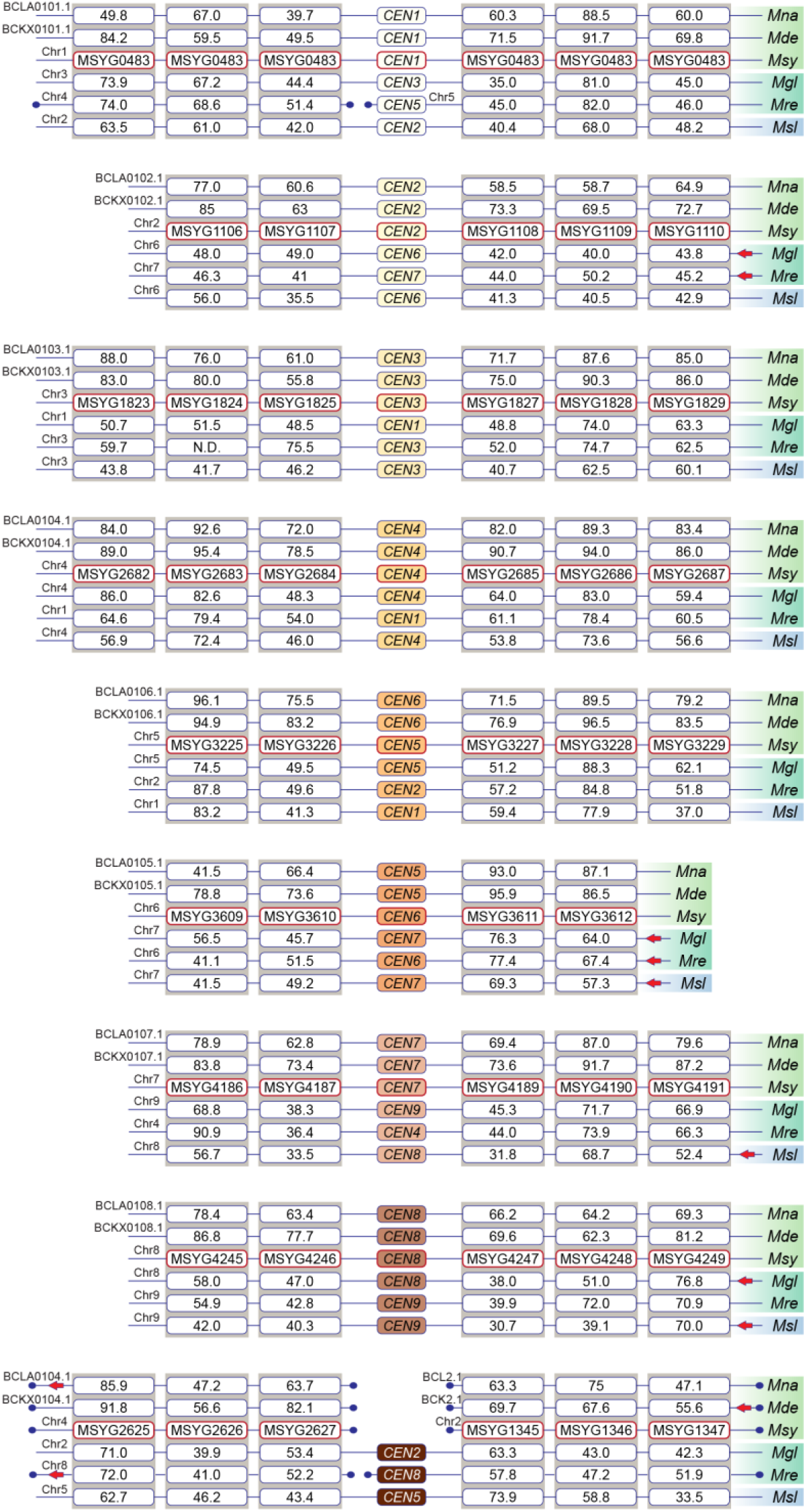
All of the predicted centromeres of *M. sympodialis, M. nana,* and *M. dermatis* belong to conserved gene synteny blocks when compared with species containing 9 chromosomes. Gene order and synteny of ORFs flanking the centromeres of 6 *Malassezia* species representative of clade B and C analyzed in this study by BLAST analysis using the protein sequences from *M. sympodialis* as query. Species included are as follows: *Mna- M. nana, Mde- M. dermatis, Msy- M. sympodialis, Mgl- M. globosa, Mre-M. restricta*, and *Msl- M. slooffiae*. Chromosome/scaffold numbers are indicated at the start of every track. Boxes represent ORFs with the numbers inside them indicating percentage identity from BLAST analysis. Broken lines marked with filled circle towards the ends indicate gene synteny breakpoints. Red arrows indicate inverted orientation of the scaffold/chromosome.

**Figure 7-figure supplement 4.**
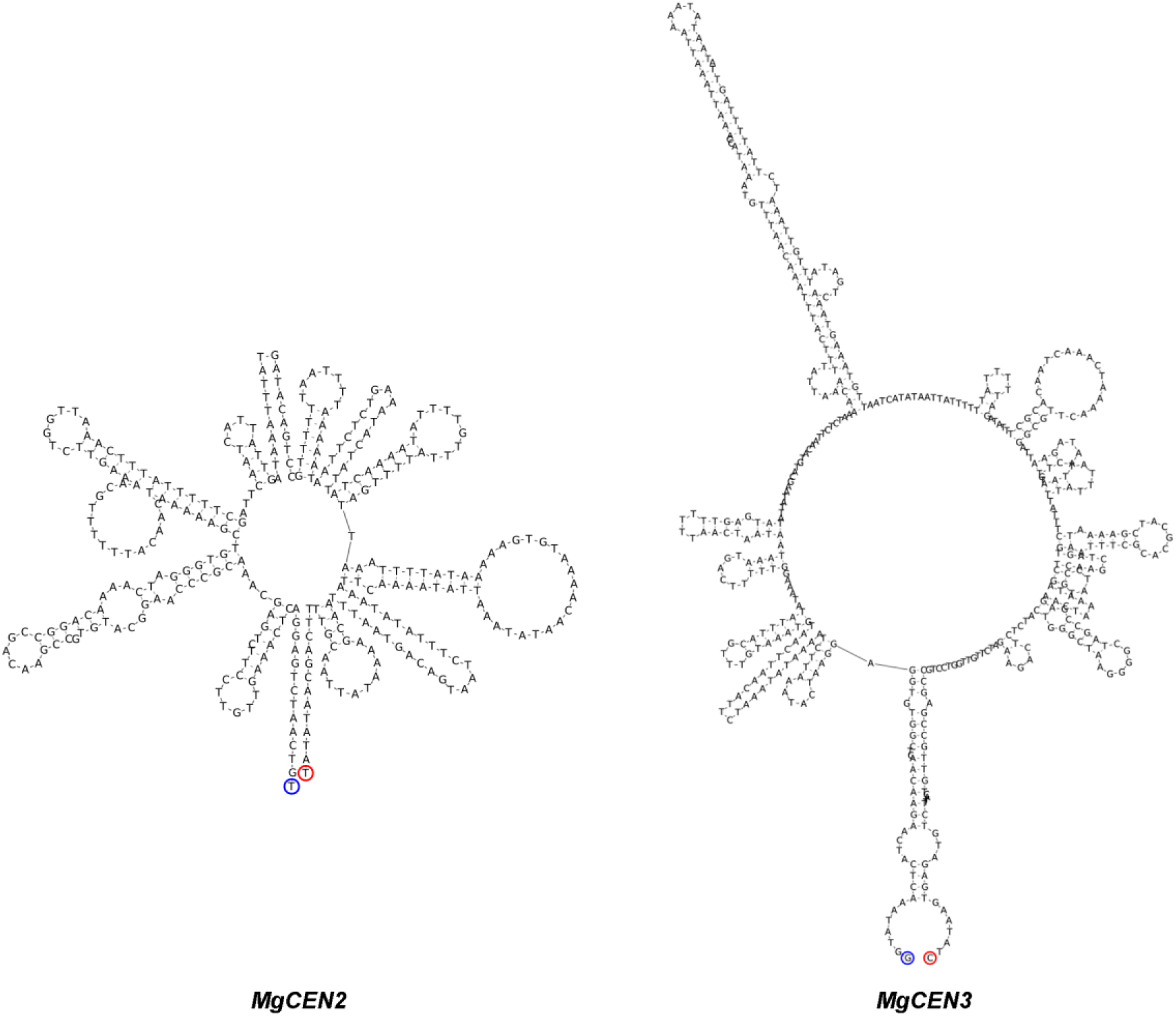
*MgCEN2* and *MgCEN3* are predicted to form secondary structures. The secondary structure of the centromere cores of *MgCEN2* and *MgCEN3* predicted by the ViennaRNA, generated using Geneious 9.0 (Lorenz et al., 2011). Blue and red circles mark the 5’- and the 3’-DNA ends respectively.

