## Supporting tables for "Loss of centromere function drives karyotype evolution in closely related *Malassezia* species"

### Supplementary tables

**Supplementary table S1. Coordinates of centromeres and their GC content in *M. sympodialis.***

Coordinates and length of Mtw1-enriched regions in comparison with that of the core centromeres in *M. sympodialis.*

| Chromosome number | Core centromere | | | | Full length centromere | | |
|---|---|---|---|---|---|---|---|
| | Coordinates | | Length (bp) | %GC | Coordinates | | Length (bp) |
| | Start | End | | | Start | End | |
| 1 | 786541 | 787061 | 520 | 16.4 | 784833 | 788599 | 3767 |
| 2 | 355760 | 355841 | 81 | 20 | 354218 | 357486 | 3269 |
| 3 | 237534 | 238686 | 1152 | 15.6 | 235615 | 239940 | 4326 |
| 4 | 418202 | 418728 | 526 | 15.2 | 415985 | 420656 | 4672 |
| 5 | 125056 | 125220 | 164 | 18 | 123219 | 127284 | 4066 |
| 6 | 101950 | 102502 | 552 | 14.4 | 100342 | 105251 | 4910 |
| 7 | 431542 | 431987 | 445 | 13.2 | 430028 | 433194 | 3167 |
| 8 | 24694 | 25564 | 870 | 18.4 | 22334 | 27476 | 5143 |

Genome average GC content (in %): 58.5

**Supplementary table S2. Coordinates of centromeres in *Malassezia* species analyzed in this study.**

Coordinates, length, and GC content (in %) of the centromeres predicted in all of the *Malassezia* species analyzed in this study.

| | Chr./Scaffold | *CEN* | Core centromere | | | | % GC Genome |
|---|---|---|---|---|---|---|---|
| | | | Start | End | Length (bp) | % GC | |
| ***M. furfur*** CBS14141 | Chr1 | *CEN1* | 2,850,135 | 2,850,402 | 268 | 15.7 | 64.9 |
| | Chr2 | *CEN2* | 68,763 | 68,931 | 168 | 15.4 | |
| | Chr3 | *CEN3* | 717,557 | 718,084 | 528 | 22.9 | |
| | Chr4 | *CEN4* | 155,897 | 156,301 | 405 | 18.3 | |
| | Chr5 | *CEN5* | 342,885 | 343,372 | 488 | 21.5 | |
| | Chr6 | *CEN6* | 86,112 | 86,832 | 721 | 27 | |
| | Chr7 | *CEN7* | 56,894 | 57,339 | 445 | 20.9 | |
| ***M. globosa*** CBS7966 | Chr1 | *CEN1* | 981894 | 982242 | 349 | 17.7 | 52.05 |
| | Chr2 | *CEN2* | 362480 | 362807 | 327 | 25.9 | |
| | Chr3 | *CEN3* | 219647 | 220121 | 474 | 27.2 | |
| | Chr4 | *CEN4* | 152635 | 152994 | 359 | 18.3 | |
| | Chr5 | *CEN5* | 215437 | 215595 | 158 | 17 | |
| | Chr6 | *CEN6* | 464007 | 464114 | 107 | 32.4 | |
| | Chr7 | *CEN7* | 736701 | 737015 | 314 | 18.1 | |
| | Chr8 | *CEN8* | 59472 | 59817 | 345 | 19.7 | |
| | Chr9 | *CEN9* | 114080 | 114535 | 455 | 23.5 | |
| ***M. slooffiae*** CBS7956 | Chr1 | *CEN1* | 138,919 | 139,465 | 547 | 26 | 66.31 |
| | Chr2 | *CEN2* | 132,717 | 133,193 | 477 | 23.1 | |
| | Chr3 | *CEN3* | 367,665 | 368,177 | 513 | 23.8 | |
| | Chr4 | *CEN4* | 130,942 | 131,501 | 560 | 27 | |
| | Chr5 | *CEN5* | 183,442 | 183,981 | 540 | 28.5 | |
| | Chr6 | *CEN6* | 411,984 | 412,552 | 569 | 27.4 | |
| | Chr7 | *CEN7* | 54,307 | 54,889 | 583 | 30 | |
| | Chr8 | *CEN8* | 497,637 | 498,149 | 513 | 24 | |
| | Chr9 | *CEN9* | 55,948 | 56,479 | 532 | 26.7 | |
| ***M. restricta*** CBS877 | Chr1 | *CEN1* | 347,813 | 348,406 | 594 | 29.7 | 55.73 |
| | Chr2 | *CEN2* | 87,190 | 87,806 | 617 | 33.3 | |
| | Chr3 | *CEN3* | 1,101,494 | 1,102,083 | 590 | 33.9 | |
| | Chr4 | *CEN4* | 754,356 | 754,989 | 634 | 34.2 | |
| | Chr5 | *CEN6* | 621,177 | 621,863 | 687 | 31.7 | |
| | Chr6 | *CEN7* | 390,657 | 391,286 | 630 | 35.1 | |
| | Chr7 | *CEN8* | 362,842 | 363,381 | 540 | 32 | |
| | Chr8 | *CEN9* | 117,021 | 117,603 | 583 | 32.8 | |
| | Chr9 | *CENR* | 70,306 | 70,913 | 608 | 36.3 | |
| ***M. nana*** JCM12085 | BCLA01000001.1 (Scf1) | *CEN1* | 715,036 | 715,592 | 557 | 27.8 | 57.95 |
| | BCLA01000002.1 (Scf2) | *CEN2* | 349,428 | 350,120 | 693 | 33 | |
| | BCLA01000003.1 (Scf3) | *CEN3* | 220,773 | 221,345 | 573 | 27.9 | |

| | | | | | | | |
|---|---|---|---|---|---|---|---|
| | BCLA01000004.1 (Scf4) | *CEN4* | 410,594 | 411,387 | 794 | 33.2 | |
| | BCLA01000005.1 (Scf5) | *CEN5* | 524,594 | 525,105 | 512 | 24.8 | |
| | BCLA01000006.1 (Scf6) | *CEN6* | 133,647 | 134,324 | 678 | 33.6 | |
| | BCLA01000007.1 (Scf7) | *CEN7* | 408,363 | 409,067 | 705 | 34.2 | |
| | BCLA01000008.1 (Scf8) | *CEN8* | 398,756 | 399,423 | 668 | 32.5 | |
| ***M. dermatis*** JCM11348 | BCKX01000001.1 (Scf1) | *CEN1* | 711,456 | 711,978 | 523 | 22.8 | 59.05 |
| | BCKX01000002.1 (Scf2) | *CEN2* | 1,014,281 | 1,014,977 | 697 | 31.7 | |
| | BCKX01000003.1 (Scf3) | *CEN3* | 232,065 | 232,795 | 731 | 29.3 | |
| | BCKX01000004.1 (Scf4) | *CEN4* | 409,839 | 410,631 | 793 | 29.5 | |
| | BCKX01000005.1 (Scf5) | *CEN5* | 94,520 | 95,018 | 499 | 18.2 | |
| | BCKX01000006.1 (Scf6) | *CEN6* | 473,487 | 474,334 | 848 | 30.4 | |
| | BCKX01000007.1 (Scf7) | *CEN7* | 76,361 | 76,975 | 615 | 26 | |
| | BCKX01000008.1 (Scf8) | *CEN8* | 17,893 | 18,540 | 648 | 26.4 | |
| ***M. vespertilionis*** CBS15041 | KZ454987.1 (Scf1) | *CEN1* | 410,820 | 411,340 | 521 | 15.7 | 56.6 |
| | KZ454988.1 (Scf2) | *CEN2* | 1,275,509 | 1,276,238 | 730 | 25.8 | |
| | KZ454989.1 (Scf3) | *CEN3* | 322,361 | 323,277 | 917 | 38.2 | |
| | KZ454990.1 (Scf4) | *CEN4* | 583,450 | 584,319 | 870 | 28.9 | |
| | KZ454991.1 (Scf5) | *CEN5* | 802,843 | 804,042 | 1,200 | 28.8 | |
| | KZ454992.1 (Scf6) | *CEN6* | 739,896 | 740,558 | 663 | 22.3 | |
| | KZ454993.1 (Scf7) | *CEN7* | 268,699 | 269,626 | 928 | 28.8 | |
| | KZ454994.1 (Scf8) | *CEN8* | 10,985 | 11,865 | 881 | 28 | |
| | KZ454995.1 (Scf9) | *CEN9* | 19,047 | 19,724 | 678 | 29.1 | |
| ***M. japonica*** JCM11963 | BCKY01000001.1 (Scf1) | *CEN1* | 1068050 | 1068614 | 564 | 25.1 | 62.35 |
| | BCKY01000002.1 (Scf2) | *CEN2* | 139423 | 139920 | 497 | 20.3 | |
| | BCKY01000003.1 (Scf3) | *CEN3* | 350068 | 350603 | 535 | 24.3 | |
| | BCKY01000004.1 (Scf4) | *CEN4* | 380877 | 381439 | 562 | 24.2 | |
| | BCKY01000005.1 (Scf5) | *CEN5* | 507632 | 508230 | 598 | 24.5 | |
| | BCKY01000006.1 (Scf6) | *CEN6* | 240968 | 250550 | 582 | 23.7 | |
| | BCKY01000007.1 (Scf7) | *CEN7* | 286711 | 287234 | 523 | 24.9 | |
| | BCKY01000008.1 (Scf8) | *CEN8* | 87314 | 87873 | 559 | 23.9 | |
| | BCKY01000010.1 (Scf10) | *CEN9* | 230906 | 231456 | 530 | 24.1 | |

**Supplementary table S3. Synteny of centromeres across all of the *Malassezia* species analyzed in this study.**

| **Clade C** | **Clade B1** | | | **Clade B2** | | **Clade A** | | |
|---|---|---|---|---|---|---|---|---|
| ***M. slooffiae*** | ***M. dermatis*** | ***M. nana*** | ***M. sympodialis*** | ***M. globosa*** | ***M. restricta*** | ***M. vespertilionis*** | ***M. japonica*** | ***M. furfur*** |
| **9 Chr** | **8 Chr** | **8 Chr** | **8 Chr** | **9 Chr** | **9 Chr** | **9 Chr** | **9 Chr** | **7 Chr** |
| *CEN2* | *CEN1* | *CEN1* | *CEN1* | *CEN3* | *CEN5* partial | *CEN6* | *CEN8* | *CEN4* |
| *CEN6* | *CEN2* | *CEN2* | *CEN2* | *CEN6* | *CEN7* | *CEN9* | *CEN4* partial | *CEN2* partial |
| *CEN3* | *CEN3* | *CEN3* | *CEN3* | *CEN1* | *CEN3* | *CEN7* partial | *CEN6* partial | *CEN3* partial |
| *CEN4* | *CEN4* | *CEN4* | *CEN4* | *CEN4* | *CEN1* | *CEN1* | *CEN2* | *CEN7* partial |
| *CEN1* | *CEN6* | *CEN6* | *CEN5* | *CEN5* | *CEN2* | *CEN4* | *CEN7* | *CEN1* |
| *CEN7* | *CEN5* | *CEN5* | *CEN6* | *CEN7* | *CEN6* | *CEN5* partial | *CEN5* partial | *CEN6* |
| *CEN8* | *CEN7* | *CEN7* | *CEN7* | *CEN9* | *CEN4* | *CEN2* | *CEN9* partial | **Inactivated** |
| *CEN9* | *CEN8* | *CEN8* | *CEN8* | *CEN8* | *CEN9* | *CEN8* | *CEN1* | **Inactivated** |
| *CEN5* | **BP** | **BP** | **BP** | *CEN2* | *CEN8* partial | *CEN3* | *CEN3* partial | *CEN5* |

'BP' indicates the presence of a gene synteny break, 'Inactivated' indicates centromere inactivation due to sequence divergence and erosion of AT-richness.

**Supplementary table S4. List of key reagents used in this study.**

| Antibodies and reagents | Source | Identifier |
|---|---|---|
| mouse anti-GFP | Roche | 11814460001 |
| mouse anti-PSTAIRE | Abcam | Cat. no.10345 |
| goat anti-mouse HRP | Bangalore Genei | Cat. no. HP06 |
| rabbit anti-H3 | Abcam | Cat. no. ab1791 |
| rabbit anti-H4 | Abcam | Cat. no. ab10158 |
| mouse anti-FLAG | Sigma | Cat. no. F1804 |
| Lysing enzymes from *Trichoderma harzianum* | Sigma | Cat. no. L1412 |
| Zymolyase 20T | MP biomedicals | Cat. no. 320921 |
| GFP-trap beads | ChromoTek | Cat. no. gta-20 |
| Blocked agarose beads | ChromoTek | Cat. no. bab-20 |
| Protein-A sepharose beads | Sigma | Cat. no. P3391 |
| M2 anti-FLAG affinity gel | Sigma | Cat. no. A2220 |
| 2-mercaptoethanol | HiMedia | Cat. no. MB041 |

**Supplementary table S5. List of strains used in this study.**

| Yeast Strains | |
|---|---|
| ATCC42132 | Wild-type *Malassezia sympodialis* |
| MSY001 | *GFP-MTW1-NAT* in *M. sympodialis* ATCC42132 |
| CBS7966 | Wild-type *Malassezia globosa* |
| CBS7956 | Wild-type *Malassezia slooffiae* |
| CBS14141 | Wild-type *Malassezia furfur* |
| MF001 | *CENPA-3xFLAG-NAT* in *M. furfur* CBS14141 |
| BY4741 | Wild-type *S. cerevisiae* strain (*MAT*a *his3Δ1 leu2Δ0 met15Δ0 ura3Δ0*) |

**Supplementary table S6. List of plasmids employed in this study.**

| **Plasmids** | |
|---|---|
| Plasmid: pGI3 | (Ianiri et al., 2017) |
| Plasmid: pVY7 | (Kozubowski et al., 2013) |
| Plasmid: pAIM1 | (Ianiri et al., 2016) |
| Plasmid: pMF1 (pGI3-MfCENP-A-3xFLAG) | This study |
| Plasmid: pMHR04 (pGI3-GFP-MsyMtw1) | This study |

**Supplementary table S7. List of oligonucleotides utilized in this study.**

| **Primers to generate epitope tagging alleles in *M. sympodialis*** | | |
|---|---|---|
| Msy Mtw1 N-P1 | GCGCGCCTAGGCCTCTGCAGGTCGACTCTGACTGACCACGACGAGCTG | Primers to tag Mtw1 with GFP at N-terminus |
| Msy Mtw1 N-P2 | CGCCCTTGCTCACCATCGAGGGGTGGAGGTACAATAG | |
| Msy Mtw1 N-P3 | CTATTGTACCTCCACCCCTCGATGGTGAGCAAGGGCG | |
| Msy Mtw1 N-P4 | GCGTCCGAGGTGGACATGTACAGCTCGTCCATGCC | |
| Msy Mtw1 N-P5 | GGCATGGACGAGCTGTAcATGTCCACCTCGGACGC | |
| Msy Mtw1 N-P6 | GAGGATCTGCACCGTGGCACATTGCGCGATGATG | |
| Msy Mtw1 N-P7 | CATCATCGCGCAATGTGCCACGGTGCAGATCCTC | |
| Msy Mtw1 N-P8 | TGATTACGAATTCTTAATTAAGATATCGAGCGTCCTCTCCTATGTCTGACC | |
| **Primers to generate epitope tagging alleles in *M. furfur*** | | |
| MfCse4 P1 | GCGCGCCTAGGCCTCTGCAGGTCGACTCTATGCAGCAACAGGCACACATG | Primers to tag CENP-A with 3xFLAG tag at C terminus |
| MfCse4 P2 | CTACTTGTCATCGTCATCCTTGTAGTCGATGTCATGATCTTTATAATCACCGTCATGGTCTTTGTAGTCCCGGATGTCGCCCCAATG | |

| | | |
|---|---|---|
| Mf NAT-F | GACTACAAAGACCATGACGGTGATTATAAAGATCATGACATCGACTACAAGGATGACGATGACAAGTAGTCCACGGTGCAGATCCTCG | |
| Mf NAT-R | GCTTTCATAGGAACATGCCCTGCGTCCTCTCCTATGTCTG | |
| MfCse4 P3 | CAGACATAGGAGAGGACGCAGGGCATGTTCCTATGAAAGC | |
| MfCse4 P4 | TGATTACGAATTCTTAATTAAGATATCGAGGAGGCGATCAACCGGCTTAG | |
| **Primers for *M. sympodialis* centromeres** | | |
| MS1 F1 | AAGAATTGATAACATTGTTGCAC | *MsyCEN1* primers |
| MS1 R1 | TAGAATAAAATGTCGCGAAGG | |
| MS2 F1 | CTGAAGAAAAGAAACAAATTCG | *MsyCEN2* primers |
| MS2 R1 | TCGGAAATCCCGCAAAAG | |
| MS3 F1 | CATATTCAGCCTCCACTAAG | *MsyCEN3* primers |
| MS3 R1 | CCTCTATCGAGTGCTCTAC | |
| MS4 F1 | CGATATGGATTGGACTTATAAGTC | *MsyCEN4* primers |
| MS4 R1 | AAAAGCAATACGTAGACGG | |
| MS5 F1 | AAATTACCGACCAGAATTG | *MsyCEN5* primers |
| MS5 R1 | ATCTGTGTCCGCTCTCATC | |
| MS6 F1 | TTTGACGCTTTATTTGTGTTTC | *MsyCEN6* primers |
| MS6 R1 | CACATATGCACGAATAATAAAACG | |
| MS7 F1 | GATACATATTCTTACACTAATACTATTCG | *MsyCEN7* primers |
| MS7 R1 | GCATAGAGCTAATATCTGATATTC | |
| MS8 F1 | GGAAGCATGAGATATTGG | *MsyCEN8* primers |
| MS8 R1 | AAACAAAGTAAAATTCTAATCACG | |
| MS8 LF1 | CTCCTCCGATACGATTCAC | *MsyCEN8* L1 primers |
| MS8 LR1 | CAGCCATTATCTCCGACAC | |

| | | |
|---|---|---|
| MS8 LF2 | CTGGGTAGATTGAGAATGAG | *MsyCEN8* L2 primers |
| MS8 LR2 | CATGTATGTTCAGTCCCATG | |
| MS8 RF1 | ATGATCCAAAAGAAAGCATAC | *MsyCEN8* R1 primers |
| MS8 RR1 | GAAGTATGTCTGGGTGAAGC | |
| MS C3 | GAAGACGACAACGATACC | Control primers away from *MsyCEN1* |
| MS C4 | TAGCGAGTGAATAGCGTC | |
| **Primers for chromoblot analysis in *M. globosa*** | | |
| Maglo_CBS7966_v2_Chr3_216001_216700_Forward: | GATGAGCGACGGAAACAAGC | Probe for Chr3 |
| Maglo_CBS7966_v2_Chr3_216001_216700_Reverse: | AACTTCGTCCCATTCGCCTT | |
| Maglo_CBS7966_v2_Chr4_150001_150700_Forward: | CATCGAGATTGCAACACAGC | Probe for Chr4 |
| Maglo_CBS7966_v2_Chr4_150001_150700_Reverse: | TGAACACAGGCGCCATTGTA | |
| Maglo_CBS7966_v2_Chr5_213001_213600_Forward: | TGCAATGAAGTCCGGCATGA | Probe for Chr5 |
| Maglo_CBS7966_v2_Chr5_213001_213600_Reverse: | AGGCACACGTTCATCTGGTT | |
| Maglo_CBS7966_v2_Chr6_461001_461600_Forward: | TGCTCACCCAAAAGACGACC | Probe for Chr6 |

| | | |
|---|---|---|
| Maglo_CBS7966_v2_Chr6_461001_461600_Reverse: | CGCGGACCTGGAACTGTATT | |
| **Primers for *M. globosa* centromeres** | | |
| Mg 1F | GAATTGCAATAGTAAGCCGAAC | *MgCEN1* |
| Mg 1R | GAATTATTCAACCCTTTGTACATC | |
| Mg 2F | GCAAAAGTTCTGGTTAAAC | *MgCEN2* |
| Mg 2R | TTCGTTAAATTACTGTCATTAG | |
| Mg 3F | GCATGTACAATTCTCTAAAAC | *MgCEN3* |
| Mg 3R | CAAGTTATCTTAATCCGCAAG | |
| Mg 4F | CAGAAAATAATAGTGATTGATAC | *MgCEN4* |
| Mg 4R | ATTTAAGATACATACACAATGC | |
| Mg 6F | GGAAATCCTGCGAGAATC | *MgCEN6* |
| Mg 6R | GCTGAATTCATAGAATCATTGAG | |
| Mg 7F | GATGATCCCAGTAACAACTG | *MgCEN7* |
| Mg 7R | GGTAGAATTGAATTTGTGTTTATC | |
| Mg 8F | GACTAGCGAATAAATCAATTGAC | *MgCEN8* |
| Mg 8R | TTAACCGTACCGAAAAACC | |
| Mg 9F | GAAAATAGTGACTGGTGGAC | *MgCEN9* |
| Mg 9R | GATTCTATTGCTATATTGTGCTTC | |
| Mg 5F | CTAAAAATGAAATTTGGGATAAAAC | *MgCEN5* |
| Mg 5R | AAGCACGATAAAAATCATAGC | |
| Mg2 L2F | CGTACCTTGTCCAAGAGC | *MgCEN2* L2 primer pair |
| Mg2 L2R | AGATCCATAGGCTTTGAATGC | |
| Mg2 L1F | ACCTTCGATTCTGTGACAC | *MgCEN2* L1 primer pair |
| Mg2 L1R | TGTTACACACTTTGCTTCGG | |

| | | |
|---|---|---|
| Mg2 R1F | AGGTCCTGACGATGTAATTG | *MgCEN2* R1 primer pair |
| Mg2 R1R | GTTGTTGATGTATGTCGTTCATG | |
| Mg2 R2F | AGCTATGCGATGTTGTTCTG | *MgCEN2* R2 primer pair |
| Mg2 R2R | CAGACGAGGAACTATTGTGAG | |
| Mg C5 | GCATAACATACGAGGATGTGC | Primer for control locus |
| Mg C6 | ATAGTGCCTGAATCTGCTG | |
| **Primers for *M. slooffiae* centromeres** | | |
| Slo1 FP | CAAATGAGCACAAACGTTG | *MslCEN1* |
| Slo1 RP | GGTAATTTACATTTCTTGTG | |
| Slo2 FP | ACTCAATAATCCAATAGAACC | *MslCEN2* |
| Slo2 RP | GAGAAAACATAAATGGTAGG | |
| Slo3 FP | AAACCGATTATCAATTCTCAAATG | *MslCEN3* |
| Slo3 RP | GTATCTGATTTGAAAACCTTCG | |
| Slo4 FP | TTCACGTGTAGCTACTTG | *MslCEN4* |
| Slo4 RP | AAATACAACAAACAACTAAAACG | |
| Slo5 FP | GAGCTGTGCAAGGTTAG | *MslCEN5* |
| Slo5 RP | GCCAAACAACGATGACG | |
| Slo6 FP | AATAGATTTGACAACCTTTGC | *MslCEN6* |
| Slo6 RP | TGCACAATTGTAAGAAAGC | |
| Slo7 FP | AATGCCAGATGATAAACTAGCTG | *MslCEN7* |
| Slo7 RP | GACTTCTGGCATAACTATTGG | |
| Slo8 FP | TTATGCTATTGTTTGAATCCG | *MslCEN8* |
| Slo8 RP | CTATCATTAAACGGAGAATACTC | |
| Slo9 FP | GACCTAGCTGTGCTTTTAG | *MslCEN9* |
| Slo9 RP | TTTCAGCAGCTTATTAGGC | |

| | | |
|---|---|---|
| Slo C1 | ACGACAAGCGTGTAAGG | Primers for control locus |
| Slo C2 | CCAACTTCTTCCTGCAG | |
| Slo1 L2F | GGAACGTGACGAGATCAC | *MslCEN1* L2 primers |
| Slo1 L2R | GTGTAGATCCGAAGTCATCAC | |
| Slo1 L1F | ATCTCTGCAAGCTTCGG | *MslCEN1* L1 primers |
| Slo1 L1R | AGTGGATGCTTCATCTTCTG | |
| Slo1 R1F | CGAATGACTTCCTCAATGC | *MslCEN1* R1 primers |
| Slo1 R1R | TGCAACAGCAGAAGAGTC | |
| Slo1 R2F | ATGCCGACCACAATCC | *MslCEN1* R2 primers |
| Slo1 R2R | ACTGTGCCTGTTTCGC | |
| **Primers for *M. furfur* centromeres** | | |
| MF1 F1 | GATAGCAAACATGATTAAAGTAATAAC | *MfCEN1* |
| MF1 R1 | GACCAAAATAATATATATTAACAAATG | |
| MF2 F1 | CAAAAGTGAAGAAGCAGG | *MfCEN2* |
| MF2 R1 | CACATATAAGAAGTAGAAAAGAAAACTC | |
| MF3 F1 | CATGTCTGGACCTCGG | *MfCEN3* |
| MF3 R1 | CGTGGTGAGAACACAAC | |
| MF4 F1 | CCTAAACTTATGAACTGTTTATTC | *MfCEN4* |
| MF4 R1 | GTTAAGTATTCCATAATGCTC | |
| MF5 F1 | CTTCTGCCATCGTTTCTC | *MfCEN5* |
| MF5 R1 | CTTGATTGTTCCTTCGTAATTAAC | |
| MF6 F2 | CATGTATGTTAAACGTCATAGTAC | *MfCEN6* |
| MF6 R2 | CGATTTGATCTATAATAACATAC | |
| MF7 F1 | GTGAAGCTATAATATTATAGAATGAG | *MfCEN7* |
| MF7 R1 | CGTTTGAATCATTATAATACTG | |

| | | |
|---|---|---|
| MF7 LF1 | GAAAGCTTCATTCGGAGC | *MfCEN7* L1 primer pair |
| MF7 LR1 | CGTCTTGGGAAGAGCAG | |
| MF7 LF2 | GGCGGATCATCTTTTCG | *MfCEN7* L2 primer pair |
| MF7 LR2 | GATTCTGATCGTCGGAGG | |
| MF7 RF1 | GTGGCACTACTGGATCG | *MfCEN7* R1 primer pair |
| MF7 RR1 | CGTGTACCGGTACATGTG | |
| MF7 RF2 | CTGTACCGCTACCTGC | *MfCEN7* R2 primer pair |
| MF7 RR2 | GTACGAATCGAGATCAACTG | |
| GI154 | GTCGGAGAAGCAGTCAATGC | Primers for control locus |
| NAT sFP | GTGCGGAGAAGGCATTGTTC | |

**Supplementary table S8. List of software and algorithms used in this study.**

| **Software and Algorithms** | | |
|---|---|---|
| Fiji | National Institute of Health | https:/fiji.sc/ |
| Photoshop CS6 | Adobe Systems | https:/www.adobe.com |
| Excel | Microsoft | https:/products.office.com/en-in/excel |
| Word | Microsoft | https:/products.office.com/en-in/word |
| Geneious 9.0 | Biomatters Ltd. | http://www.geneious.com/ |
| SyMap | (Soderlund et al., 2011) | |
| PhylloGibbs-MP | (Siddharthan, 2008) | |
| Circos | (Krzywinski et al., 2009) | |
| Satsuma | (Grabherr et al., 2010) | |
| Easyfig | (Sullivan et al., 2011) | |
